# Structural Insights into GluK3-kainate Receptor Desensitization and Recovery

**DOI:** 10.1101/525154

**Authors:** Jyoti Kumari, Rajesh Vinnakota, Janesh Kumar

**Author notes:** **Corresponding author**: Janesh Kumar Ph.D., National Centre for Cell Science, NCCS Complex, S. P. Pune University Campus, Ganeshkhind, Pune-411 007, India.

## Abstract

GluK3-kainate receptors are atypical members of the iGluR family that reside at both the pre- and postsynapse and play key role in regulation of synaptic transmission. For better understanding of structural changes that underlie receptor recovery from desensitized state, GluK3 receptors were trapped in desensitized and resting/closed states and structures analyzed using single particle cryo-electron microscopy. We show that receptor recovery from desensitization requires major rearrangements of the ligand binding domains (LBD) while the amino terminal (ATD) and transmembrane domains remain virtually unaltered. While, the desensitized GluK3 has domain organization as seen earlier for another kainate receptor-GluK2, antagonist bound GluK3 trapped a partially “recovered” state with only two LBD domains in dimeric arrangement necessary for receptor activation. Using these structures as guide, we show that the N-linked glycans at the interface of GluK3 ATD and LBD likely mediate inter-domain interactions and attune receptor-gating properties. Mutational analysis also identifies putative N-glycan interacting residues. These results provide a molecular framework for understanding gating properties unique to GluK3 and identify role of N-linked glycosylation in their modulation.

---

Ionotropic glutamate receptors (iGluRs) mediate majority of fast excitatory neurotransmission at the chemical synapses in the central nervous system (CNS). Due to this pivotal role, their dysfunction is implicated in a wide range of neurological disorders such as schizophrenia, neuro-excitatory disorders, epilepsy, neurodegenerative, developmental disorders, neuropathic pain etc^1^. The iGluR family is broadly classified into 4 classes such as α-amino-3-hydroxy-5-methyl-4-isoxazolepropionic acid receptor (AMPA), kainate (KA), N-methyl-D-aspartate (NMDA) and orphan delta receptors. AMPA, KA and NMDA receptors form tetrameric cation conductive channels that are activated by presynaptically released glutamate resulting in influx of sodium and calcium ions leading to membrane depolarization^2^. Although, all the iGluRs share similar domain organization, membrane topology and sequence similarity, they have distinct pharmacological, physiological and biophysical properties^3^. In contrast to the AMPA and NMDA receptors that are primarily localized on post-synaptic membrane and are directly involved in neurotransmission, increasing evidence has shown that KARs are expressed at both the presynaptic and postsynaptic sites. KA receptors play a key role in regulation of synaptic networks by modulating the release of neurotransmitters at the presynaptic sites and participating in membrane depolarization at postsynapses^4^. KARs are abundantly expressed in hippocampus and cerebellum; display characteristic slow kinetics, are mainly involved in long-term memory formation and motor control^5, 6^. They are divided into two families consisting of the “low-affinity” glutamate binding subunits (GluK1-GluK3) that form functional homomeric ion channels and “high-affinity” subunits (GluK4 and GluK5) that form functional receptors only on assembly with “low-affinity” subunits^7-9^.

Of all the KARs, GluK3 is one of the least studied subunit with respect to structure-function analysis. However, investigations on GluK3 knockout mice have clearly shown their important role at presynaptic sites in the hippocampal mossy fiber synapse^10^. Electrophysiological assays have shown that unlike homomeric GluK2 or heteromeric GluK2/K5 receptors, the desensitization rates of partially bound homomeric GluK3 receptors are much faster than those of fully bound states^11^. Further, GluK3 receptors are activated by higher glutamate concentrations compared to other KARs and are potentiated by zinc^12^. Although, crystal structures and biophysical analysis of ligand binding domain (LBD) ^12, 13^ and amino terminal domain (ATD) ^14, 15^ of GluK3 receptors have been reported, full-length structures are still elusive. Hence, insights into conformational changes underlying GluK3 receptor functions in the context of full-length tetrameric receptors are lacking. Further, iGluRs including GluK3 are extensively N-glycosylated which likely imparts additional level of regulation of their functions. These glycans have been reported to affect the trafficking and gating properties of iGluRs^16-18^ however the mechanism of this regulation is unclear. The polar, charged and complex sugar chains present in N-glycans may interact with other synaptic proteins or lay in close proximity to the functional domain of iGluRs^19^ thereby affecting receptor functionality such as deactivation, desensitization and recovery from desensitization^20^. N-glycans have also been shown to be important for efficient assembly and trafficking of AMPA^21, 22^ NMDA^23, 24^ and KA receptors^20^.

In this study, we expressed and purified intact homotetrameric rat GluK3 receptors from baculovirus infected HEK293 GnTI^-^ cells, trapped them in agonist and antagonist bound states and determined their structures using single particle cryo-electron microscopy. We have also explored the involvement of N-glycosylation sites in the receptor function using structural information of GluK3 and electrophysiology. These data shed light upon molecular mechanism of the transition between these two states during the gating cycle of receptor and the importance of N-glycosylation for GluK3 function.

## Results

### Receptor purification and structure determination

The wild type GluK3 receptor was optimized to improve its expression and stability in heterologous expression system as elucidated in Fig. 1a and **Supplementary Fig.1.** The final construct referred to as GluK3_EM_ (Fig. 1a) henceforth was optimized using the fluorescence-detection size exclusion chromatography (FSEC) ^25^ and had a symmetrical profile corresponding to tetramer on size exclusion chromatography (Fig. 1b). GluK3_EM_ exhibited gating profile similar to wild type (WT) GluK3 receptors as validated by whole-cell patch clamp recordings in HEK293T cells. It had similar rectification properties, rates of entry into desensitization and recovery from desensitized state (Fig. 1c-e, Table 1). Our electrophysiological recordings are also consistent with previous reports^11^.

**Table 1:** Electrophysiological assays for the wild type and mutant GluK3 receptors

| N-Glycan knockout | Mutation Name | Desensitization (Tau2, ms) | Dose response (EC 50) mM (95% CI) | Recovery time constant (S) (95% CI) | Rise Time (ms) |
| --- | --- | --- | --- | --- | --- |
|  | **GluK3** | 1.6±0.1 (n=6) | 9.5 (n=7)<br>(8.4 to 10.7) | 0.9 (n=5)<br>(0.8 to 1.2) | 1.3±0.1 (n=7) |
|  | **GluK3<sub>EM</sub>** | 1.2±0.2 (n=6)<br>P=0.0878 |  | 1.1 (n=5)<br>(0.6 to 2.4) |  |
| **GluK3 N-Glycan Mutations** |  |  |  |  |  |
| N 247 | **T249A** | 1.8±0.3 (n=4)<br>P=0.5536 | 7.2 (n=5)<br>(6.5 to 7.9) | 0.5 (n=5)<br>(0.3 to 1.0) | 1.3±0.1 (n=5) |
| N 395 | **T397A** | 1.6±0.1 (n=10)<br>P=0.8105 | 3.7 (n=5)<br>(3.7 to 7.2) | 0.8 (n=5)<br>(0.4 to 1.9) | 1.1±0.1 n=6) |
| N 402 | **S404A** | 2.0±0.3 (n=6)<br>P=0.1988 | 10.3 (n=5)<br>(9.3 to 11.4) | 0.5 (n=4)<br>(0.4 to 0.7) | 1.4±0.1 (n=6) |
| N247/N395 | **T249A/T397A** | 1.9±0.2 (n=11)<br>P=0.3122 | 6.5 (n=4)<br>(6.0 to 7.0) | 0.6(n=6)<br>(0.4 to 0.8) | 1.7±0.1 (n=6) |
| N247/N402 | **T249A/S404A** | 2.1±0.2 (n=5)<br>*P=0.0383 | 5.3 (n=5)<br>(4.9 to 5.6) | 0.3 (n=5)<br>(0.2 to 0.4) |  |
| N395/N402 | **T397A/S404A** | 2.1±0.2 (n=9)<br>P=0.0988 | 5.7 (n=5)<br>(5.3 to 6.1) | 0.5 (n=6)<br>(0.4 to 0.6) | 1.2±0.1 (n=6) |
| **Potential N-Glycan interacting residues** |  |  |  |  |  |
|  | **E239L** | 2.0±0.2 (n=5)<br>P=0.0951 |  | 0.3 (n=4)<br>(0.2 to 0.5) | 1.6±0.1 (n=5) |
|  | **R242E** | 2.1±0.1 (n=6)<br>*P=0.0084 |  | 0.8 (n=5)<br>(0.5 to 1.6) |  |
|  | **Y243F** | 2.3±0.5 (n=5)<br>P=0.2032 |  | 0.6 (n=4)<br>(0.5 to 0.8) | 1.4±0.1 (n=5) |
|  | **R242E/Y243F** | 3.3±0.5 (n=5)<br>*P=0.0061 |  | 4.0 (n=5)<br>(2.5 to 7.2) | 1.5±0.3 (n=4) |
| **Desensitization ring residues** |  |  |  |  |  |
|  | **D672R** | 1.7±0.2 (n=5)<br>P=0.5471 |  | 0.7 (n=5)<br>(0.6 to 0.9) | 1.1±0.2 (n=6) |
| Note P values less than 0.05 are considered as of statistical significance and are marked with *. All values are rounded off to single decimal place. |  |  |  |  |  |

**Fig. 1.**
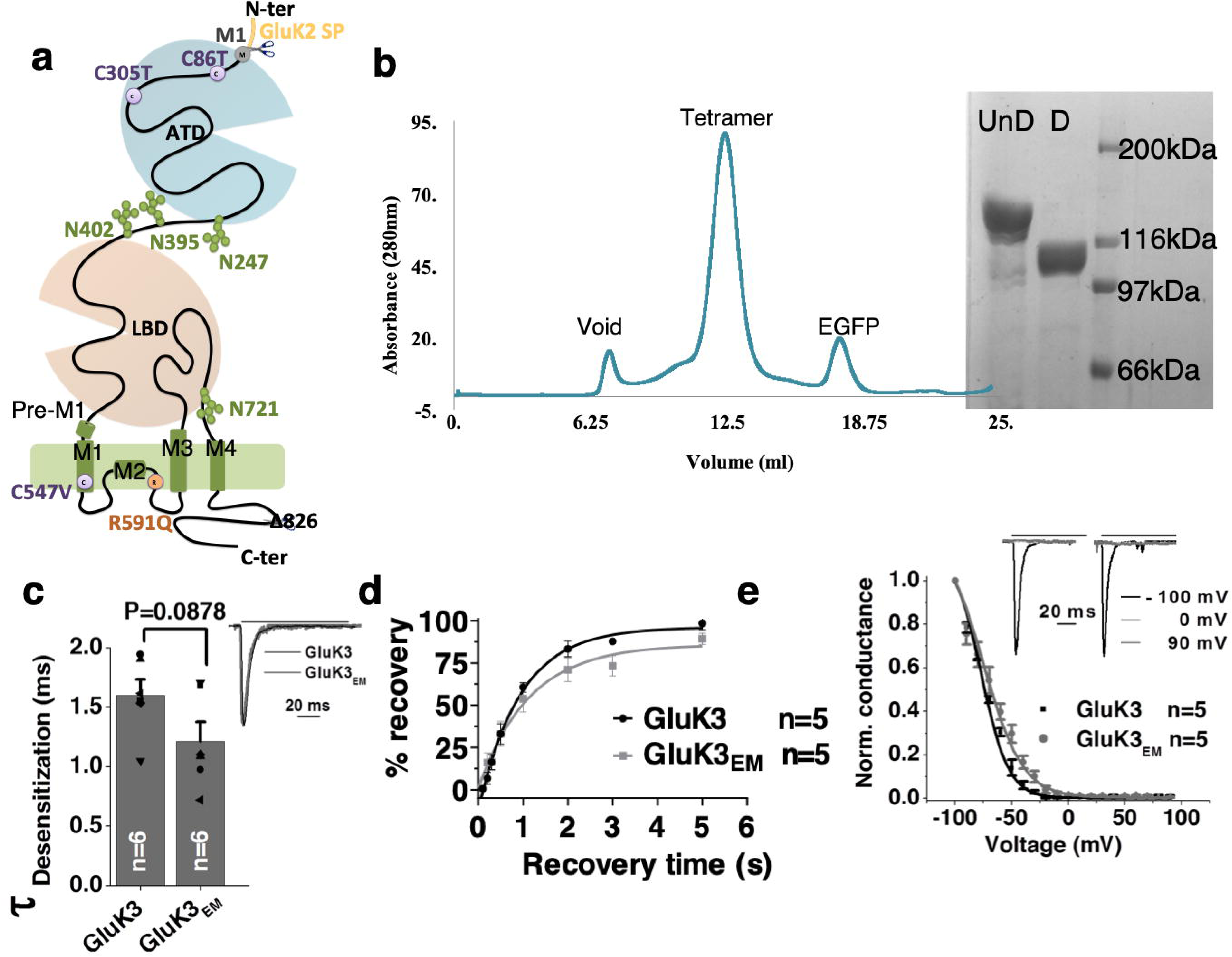
Construct design and optimization of GluK3 receptors for structural studies. **a** Schematic representation of GluK3EM construct. Amino terminal domain (ATD), ligand binding domain (LBD), and transmembrane domain (TMD) are shown alongwith the mutations made for construct optimization. GluK2 signal peptide instead of native GluK3 signal peptide and C-terminal truncation at residue 826 led to improved protein expression. Point mutations of free cysteines C86T, C305T in ATD and C547V in TM domain are shown in violet while R591Q mutant in TM is highlighted in orange. N-glycans at the ATD-LBD interface and the LBD-TM interface are also shown. **b** The SEC profile demonstrates monodisperse tetrameric receptor peak eluted from Superose 6 column and SDS-PAGE analysis of undigested (UnD) and thrombin digested (D) GluK3_EM_ is shown. Panels **c-e** show electrophysiological studies of the wild type and GluK3_EM_ construct with very similar rates for desensitization (**c**); recovery from desensitization (**d**) and indistinguishable normalized conductance measured from −100 to +100 mVolts (**e**). Representative rectification index at −100mV, 0mV and 90mV are shown for wild-type (black) and GluK3_EM_ (grey). Amplitudes of normalized steady-state currents for wild-type and GluK3_EM_ on application of 30 mM glutamate is shown in inset **c**.

In order to understand the structural basis of the receptor rearrangements during its gating cycle, we elucidated the GluK3_EM_ structure in desensitized and closed state using single particle cryo-EM. These states were captured in presence of agonist 2S, 4R-4-methylglutamate (SYM)^26^ and antagonist UBP310^27^ respectively (**Supplementary Fig. 2**). Reference-free 2D classification obtained after cryo-EM data processing for both the complexes showed good distribution of different receptor orientations as well as identifiable external features resembling glutamate receptor (**Supplementary Fig. 3a and b**). Various steps of EM data processing and 3D reconstruction of the complexes reveals similar receptor architecture as previously observed for GluA2^28, 29^ and GluK2^30, 31^ (Fig. 2; **Supplementary Fig. 4a and b**). The final density map of the closed and desensitized states has an estimated resolution of ∼7.7 Å and 7.4 Å as per 0.143 FSC gold standard respectively (**Supplementary Fig. 5 and 6**). Details of cryo-EM data collection and refinement are shown in Table 2. Our map shows well-defined features for the ATD and LBD domains while the S1-M1 and M3-S2 linker are poorly resolved (**Supplementary Fig. 7**). Co-ordinates of previously reported GluK3 ATD (PDB ID: 3OLZ) ^14^ and LBD-SYM complex structure determined in this study were fitted into EM density map for generating model of tetrameric receptor. For TM domain, a model was generated by threading GluK3 sequence onto GluK2 TM domain (PDB ID: 5KUF) ^31^. For GluK3-UBP310, an extended LBD model was generated by threading GluK3 sequence onto GluK2 LBD-LY complex (PDB ID: 5CMK) ^31^. These individual models were rigid body fitted into EM maps of GluK3-SYM and GluK3-UBP310 to generate tetrameric models in UCSF-Chimera^32^ and subsequently subjected to several rounds of real space refinement as implemented in Phenix software suite^33^.

**Table 2.** Cryo-EM data collection, refinement and validation statistics

|  | **GluK3-2S,4R-4-methylglutamate** | **GluK3-UBP310** |
| --- | --- | --- |
| **Data collection and processing** |  |  |
| Microscope | Titan Krios | Titan Krios |
| Voltage (kV) | 300 | 300 |
| No. of Micrographs | 719 | 1693 |
| Camera | Falcon 3 | Falcon 3 |
| Mode of recording | Super-resolution counting | Super-resolution counting |
| Exposure time (seconds) | 60 | 60 |
| Electron exposure (e-/Å<sup>2</sup>) | 16.73 | 17.27 |
| Defocus range (μm) | 1.5 – 3.3 | 1.5 – 3.3 |
| Pixel size (Å) | 1.38 | 1.41 |
| Symmetry | C1 | C1 |
| Initial particle images (no.) | 77,276 | 437,986 |
| Final particle images (no.) | 9,730 | 27,194 |
| Map resolution (Å) | 7.4 | 7.6 |
| FSC threshold | 0.143 | 0.143 |
| **Refinement** |  |  |
| Initial model used (PDB code) | 3OLZ (ATD),<br>XXXX(LBD), 5KUF(TM) | 3OLZ (ATD), 5CMK(LBD),<br>5KUF(TM) |
| Model resolution (Å) | 7.58 | 8.07 |
| FSC threshold | 0.5 | 0.5 |
| Map-to model fit, CC_mask | 0.8204 | 0.8248 |
| **Model composition** |  |  |
| Non-hydrogen atoms | 22889 | 23013 |
| Protein residues | 2881 | 2868 |
| **R.m.s. deviations** |  |  |
| Bond lengths (Å) | 0.007 | 0.007 |
| Bond angles (°) | 1.03 | 0.920 |
| **Validation** |  |  |
| MolProbity score | 1.99 | 1.82 |
| Clashscore | 11.57 | 12.42 |
| Ramachandran plot |  |  |
| Favored (%) | 93.9 | 96.6 |
| Allowed (%) | 6.1 | 3.4 |
| Disallowed (%) | 0.0 | 0.0 |

**Fig. 2.**
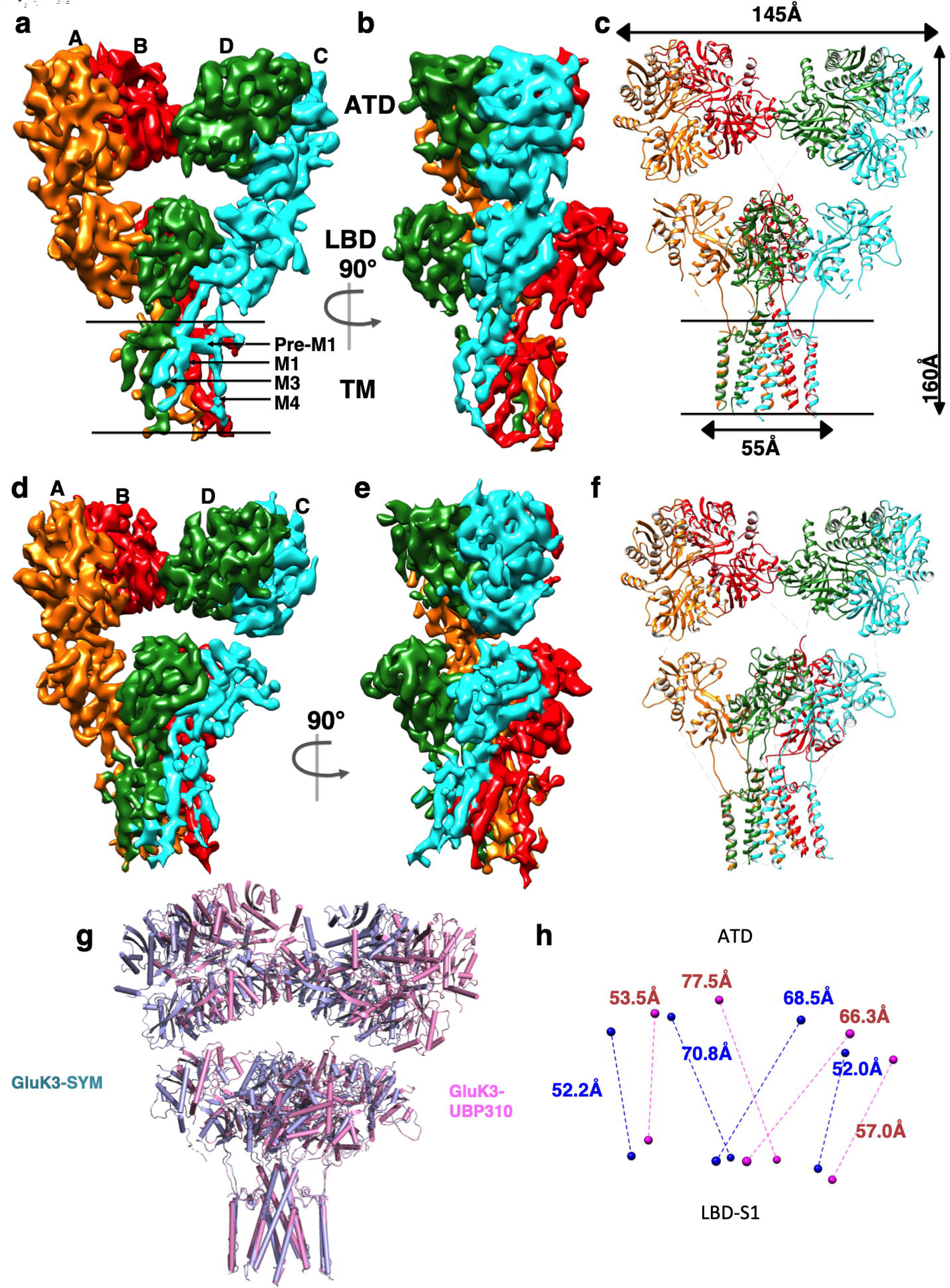
Cryo-EM density map and fitted model for the agonist and antagonist bound GluK3. Segmented and colored cryo-EM density map at 7.4 Å for agonist 2s,4R-4-methylglutamate-bound (**a-c)** and at 7.7 Å for antagonist UBP310 bound GluK3_EM_ **(d-f)**, **a** and **d** show the front view of receptor, perpendicular to the overall two-fold axis of molecular symmetry with each subunit colored uniquely; **b and e** are 90° rotated view of **a** and **b**; **c** and **f** show the fitted atomic model colored to show four receptor subunits as in **a-b and d-e**. Panel **g-h** show reorientation of the extracellular domains during transition from agonist-bound to antagonist bound state. **g** shows front view (perpendicular to the global 2-fold axis of symmetry) of the full-length GluK3 receptor structure in complex with SYM (in cyan), superposed on the UBP310 structure (magenta) by aligning the TMD regions. Distances between center of masses, shown as spheres between ATD and LBD-S1 lobe of each subunit in SYM (blue) and UBP310 (red) bound receptor is measured and indicated in panel **h**.

**Fig. 3.**
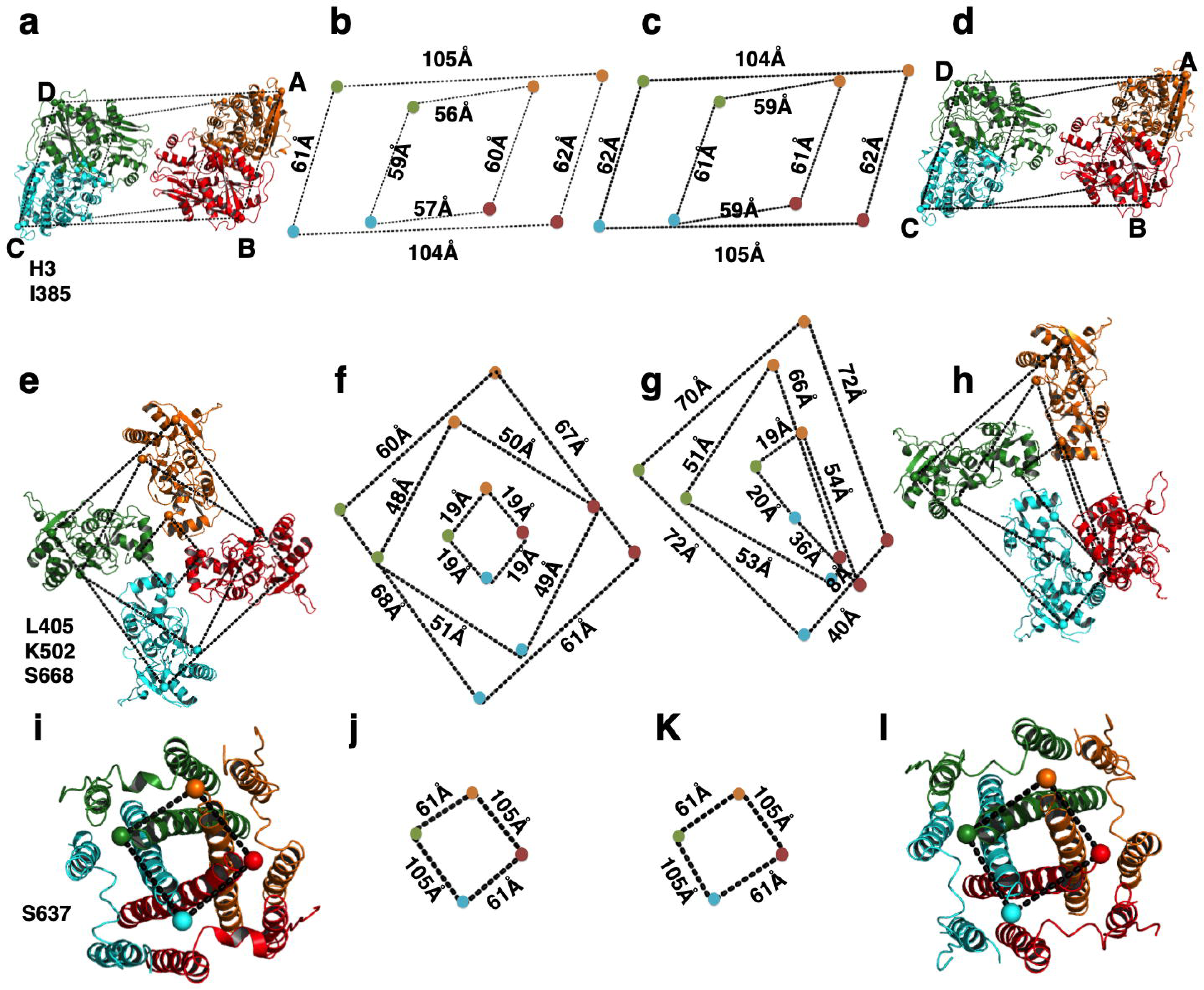
Structural comparison of extracellular and TM domains in SYM and UBP310 bound GluK3_EM_. **a-d** Top views of ATD (upper panel); **e-h** LBD (middle panel) and **i-l** TMD layers (lower panel) are shown. Cα - Cα distances between subunits for selected positions are measured and indicated. Colored dots connected by dashed lines identify the locations of His 3 and Ile 382 at the top and base of the ATD and indicate very similar parallelograms formed for both SYM (**a-b**) and UBP310 (**c-d**) bound forms. Panels **e-h** show positions of Cα atoms for Leu 405, Lys 502 and Ser 668 indicating the top, middle and base of the LBD connected by dashed lines. Symmetric parallelograms are formed at all levels of LBD for SYM **e-f**, while asymmetric trapezoids are observed in UBP310 bound state shown in **g** and **h**. Top view of the TM domains with Ser 637 connected by dashed lines is shown in **i-j** (SYM) and **k-l** (UBP310).

**Fig. 4:**
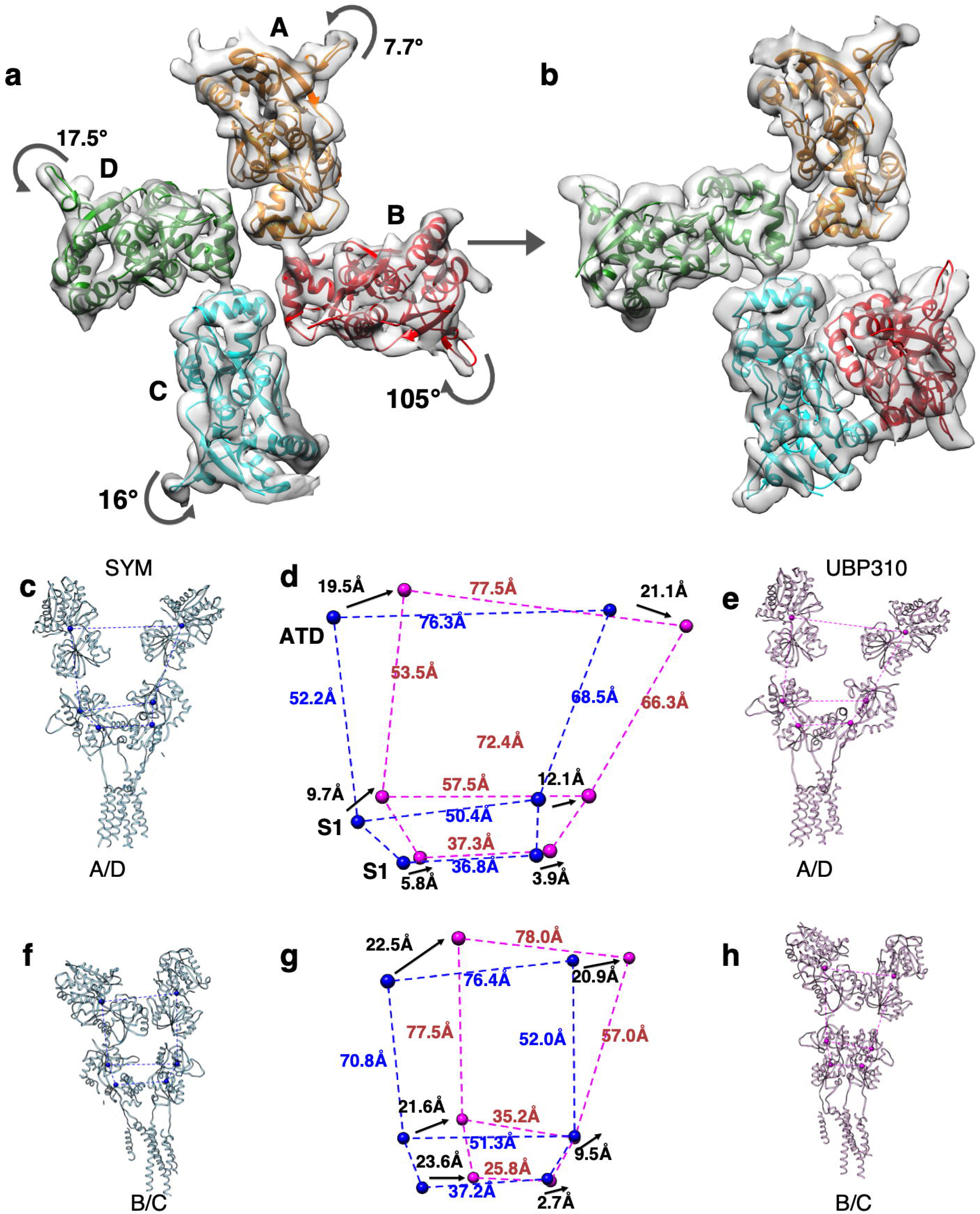
Conformational changes during transition from desensitized to resting state. **a-b**, Top views of the segmented density maps for LBD fitted with coordinates shown in ribbon in **a** agonist-bound desensitized and **b** antagonist-bound resting state. Structural changes in an LBD tetramer assembly underlying the transition from desensitized to resting state are depicted. Degree and direction of LBD rotations are measured and indicated. Panel **c-h** show subunits A/D and B/C for SYM and UBP310 bound receptor aligned on TM domains. Distances from the center of mass (spheres) between subunits for the ATD (top) and LBD S1 & S2 lobes are measured and indicated by dashed lines. Structures were superimposed using main-chain atoms of the TM domains. **d & g** show superimpositions of the parallelograms formed for the A/D (**d**) and B/C (**h**) subunits. The movements of the COM for domains between SYM and UBP310 structures are measured and indicated in black fonts, while the distances within structures are indicated in blue (SYM) and red (UBP310).

**Fig. 5.**
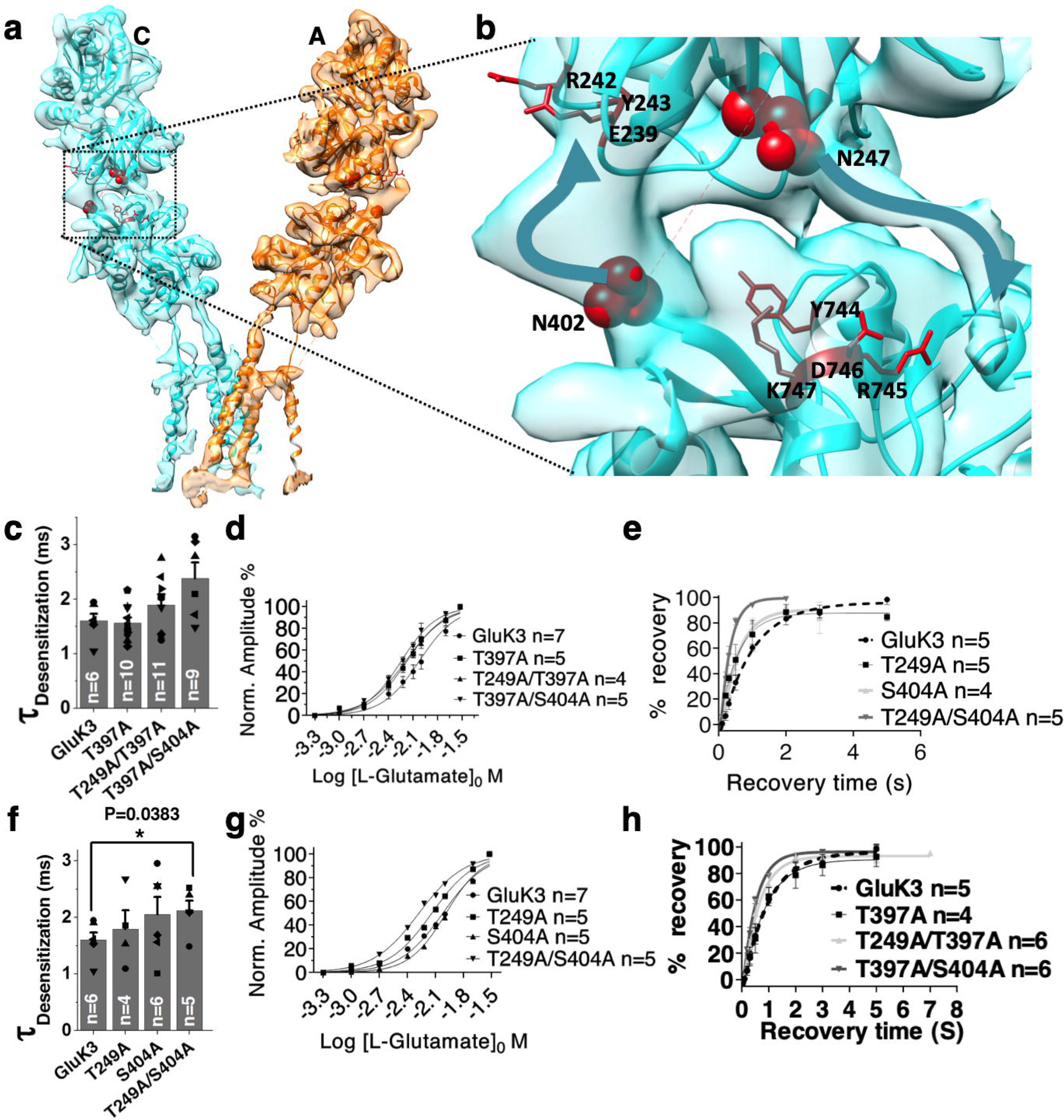
N-glycans modulate GluK3 gating properties. Panel **a** depicts the segmented density map for proximal subunits A and C fitted with atomic model. Zoomed view of the residual density at the ATD-LBD interface for subunit A (marked by black dashed lines) is shown in **b** where sites for N-glycosylation at Asn 247 and 402 are shown in red spheres. Potential interacting residues that lie in close proximity to N-glycans are shown in red stick representation. Thick blue arrow highlights the residual density. **c-h** show electrophysiological characterization of various N-glycan knockouts. Panels **c & f** show desensitization rate (τ_des_) for the single and double glycans mutants as indicated. Tau 2 values obtained from a bi-exponential fit for curve obtained by 100 ms application of 30 mM glutamate. Panels **d & g** show concentration response curves evoked by 100 ms application of glutamate at concentrations ranging from 30 µM to 30 mM normalized to the 30 mM glutamate evoked current amplitude. Panels **e & h** show two-pulse glutamate (30 mM) recovery experiments for the indicated mutants and wild type GluK3 receptors recombinantly expressed in HEK293 cells. Amplitude of the second glutamate application in a two-pulse experiment is reported as a normalized percentage of the first glutamate application and is plotted against interpulse intervals. Recovery rates (τ_rec_) were calculated with single exponential association fits.

**Fig. 6.**
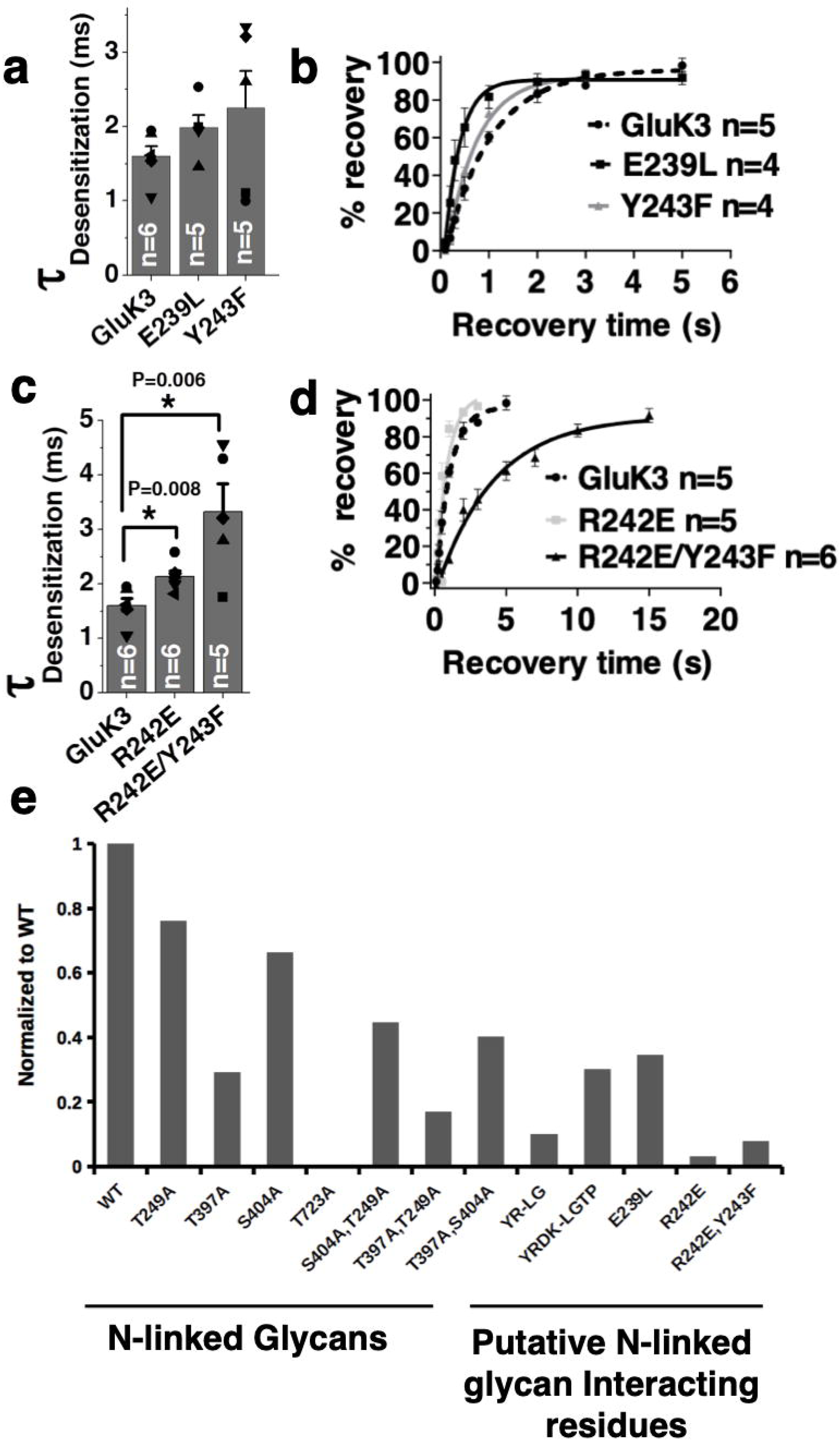
Mutations of the potential N-glycan interacting residues affect the gating properties of GluK3 receptors. Panels **a-d** show that mutations of putative interacting residues involved in inter-domain N-glycan mediated interaction affect not only desensitization kinetics (**a, c**) but also rate of recovery from the desensitized state (**b-d**). Disrupting these interactions slows the rate of receptor desensitization and hastens the recovery of receptor from desensitized state. Panel **e** shows total surface expression (in HEK293T cells) of the mutant receptors generated in this study normalized to surface expression of wild-type GluK3 as determined by surface biotinylation assay. Total and surface expressed proteins were analyzed by immunoblotting using GluR6/7 monoclonal antibody and quantified using imageJ.

**Fig. 7.**
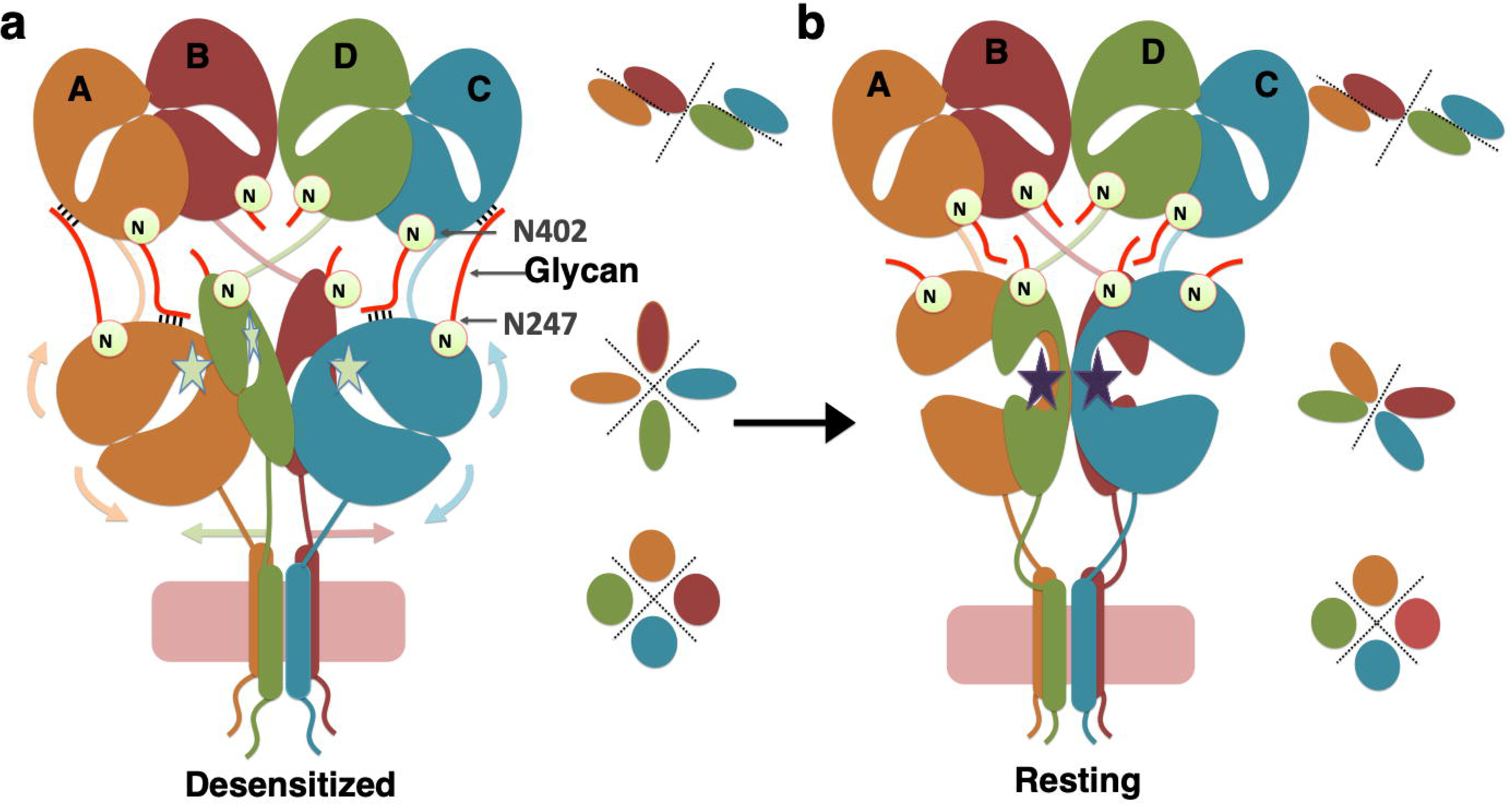
Model depicting effect of N-glycans on the receptor recovery from desensitized state *via* interdomain interactions at the ATD-LBD interface. Inter-domain interactions at the ATD-LBD interface mediated by N-glycans (thick red lines) at Asn 247 and Asn 402 positions are shown in a desensitized GluK3 receptor in panel **a**. Panel **b** shows a schematic for “fully-recovered” receptor where the N-glycan mediated interactions (depicted in black lines) would be broken due to rearrangements at the LBD layer. These interactions likely contribute in impeded receptor recovery from desensitized state since the desensitized-resting state transition requires that LBD domains return to dimeric configuration. These transitions would restore 2-2-4 symmetry for ATD, LBD and TM domains respectively in closed/resting state (**b**) from 2-4-4 symmetry observed in desensitized state (**a**) shown in cartoon for various domains.

### Desensitized and closed state GluK3 receptor structure

The desensitized and closed-state structures have a similar three-layered assembly for the ATD, LBD and TM domains as reported earlier for other iGluRs. Receptor assembly is mainly mediated by ATD as dimer of dimer (AB and CD) and then undergoes domain swapping at the LBD layer (BC and AD) as observed for AMPA^28, 34, 29^ and kainate^30, 31^ receptors.

The arrangement of ATD and TM domains in both GluK3-SYM and GluK3-UBP310 complex is similar conformation with an apparent 2-fold symmetry at dimer and dimer of dimers interface of ATD resulting into N-shaped arrangement; and 4-fold symmetry of the TM domains (Fig. 2 and 3a-d; **Supplementary Fig. 8**). This infers that the dimer of dimer at the ATD level is intact in both resting and closed state. The recently reported O-shaped arrangement of the ATD layer was not observed in our study^35^. Consistent with this, the parallelogram formed by joining Cα atoms of His 3 from each subunit at top and Ile 385 at the base has similar dimension in desensitized and closed state (Fig. 3b and c). This also suggests that there are no major structural changes at the ATD layer in order to undergo transition from desensitized to closed state. The resolution of our EM maps are adequate to (**Supplementary Fig. 7**) show unambiguously that in both desensitized and closed state the ion channel adopts a similar closed pore conformation, where the M3 helices are arranged in a crossed bundle assembly with the pre-M1 helices wrapped around the channel. Thus, the TM domains in both cases form a similar parallelogram with dimensions of ∼61 Å × 105 Å on joining Cα atoms of Ser 637 from all the subunits (Fig. 3i-l).

In contrast to ATD and TM domains, comparison of the SYM and UBP310 bound structures show major conformational changes at the LBD layer. LBD in GluK3-SYM bound state adopts an apparent 4-fold symmetry (Fig. 3e) and is similar to that reported for GluK2 receptors^30, 31^ (**Supplementary Fig. 9**). Thus, the progressively reducing parallelograms formed by joining Cα atoms of Leu 405, Lys 502 and Ser 668 at top, middle and at the base of LBD have dimensions indicative of a symmetric arrangement of both upper and lower LBD lobes (Fig. 3e-f). In the GluK3-UBP310 complex however, we observe previously unseen asymmetric arrangement for kainate receptors trapped in a putative closed state. Interestingly, subunits B and C form the LBD dimer characteristic of an antagonist-bound form seen in both GluA2 and GluK2 receptors; while the subunits A and D are separated and exist in a desensitized-like state (Fig. 3h). Due to such arrangement, joining of Cα atoms of Leu 405, Lys 502 and Ser 668 at top, middle and at the base of LBD forms an asymmetric trapezium of reducing dimensions shown in Fig. 3g-h. It is interesting to note that all the four LBD domains are in an extended cleft conformation characteristic of an antagonist-bound state in UBP310 complex; whereas SYM bound density map reveals LBD ‘clamshell’ closure (Fig. 3d **and Supplementary Fig. 10**). It is possible that we have trapped an intermediate between desensitized and “fully-recovered” state. Since, our cryo-EM analysis does not reveal any 2D or 3D class similar to “fully-recovered” state observed in GluK2 and GluA2, where both LBDs are arranged as two dimers with 2-fold symmetry, the trapped structure likely represents a stable conformation. In addition, the LBD arrangement in GluK3-UPB310 complex is similar to that observed in GluA2 complex with partial agonist fluorowillardine^29^. Hence, it is likely that the resting state in GluK3 with both the LBD dimers recovered may not be as stable compared to other AMPA and kainate receptors of known structures. It is also important to note that we have imaged the receptors saturated with ∼100 fold excess concentration of UBP310 than its EC_50_ value calculated via electrophysiology^27^. It is likely that this partial recovery contributes to low glutamate sensitivity of GluK3.

In both desensitized and closed/resting state, the ligand binding is uncoupled from the transmembrane domain as strain on the linkers connecting LBD to TM domains are relaxed by extended conformation of antagonist bound LBD (even while they are in dimeric configuration). In desensitized state, the LBD-TM linker strain due to bound agonist is relaxed via rearrangements of LBD from a dimer of dimer scheme to a pesudo-4 fold symmetric arrangement, where the LBD dimers are disrupted (Fig. 2). In concordance with this, the TM domains in both the structures adopt similar closed pore configuration.

### Transitions from desensitized to resting/closed state require large conformational changes in LBD and the LBD-TM linkers

In the gating cycle, post-desensitization, receptor needs to recover to a closed/resting state, where the LBDs couple as 2-dimers for the next cycle of activation. Hence, in a “fully-recovered” receptor, the four LBDs should rearrange into two dimers as observed previously in GluK2^31^ and GluA2^28, 29^ closed-state structures to enable coupling of agonist binding with channel opening^36^. In order to understand this transition, we compared SYM and UBP310 bound structures. It was observed that there is very little change in the conformations of the ATD and TM domains. On the contrary, major changes take place in the LBD layer and in the linkers connecting S1-M1, M3-S2 and S2-M4. Surprisingly, ligand binding domains of only subunits B and C return to a dimeric state, while subunits A and D still remain in a desensitized–like state (Fig. 4). The LBD-distal subunit B swings clockwise by ∼105° in the horizontal plane, while the LBD-proximal subunit C rotates anticlockwise by ∼16° to achieve dimeric configuration (Fig. 4a-b). In contrast to this, the LBD of subunit A and D undergo smaller degree of anti-clockwise rotation by ∼7.7° and ∼17.5° respectively and hence are unable to achieve a dimeric configuration (Fig. 4a-b). In order to further analyze structural changes in detail, we aligned both receptors at TM domains and calculated center of masses (COM) for ATD and S1; S2 lobes of LBDs to measure their displacement in going from SYM to UBP310 bound state. We observe that receptor subunits A and D swing laterally by ∼19.5 – 21 Å at the ATD level and ∼9.7 – 12 Å at the LBD-S1 to accommodate this asymmetric arrangement of LBD in UBP310 bound state. While smaller movements of ∼3.9 to 5.8 Å, are observed in S2 lobe (Fig. 4c - e). On the other hand, subunits B/C that go from monomeric to dimeric LBD configuration swing laterally by ∼20.9 - 22.5 Å at the ATD level. Interestingly, the separation between COMs for the B and C ATD remains unchanged between SYM and UBP310 bound forms suggesting a rigid body movement of the entire ATD layer (Fig. 4f - h). The movements are more pronounced and asymmetric in the LBD layer for B/C subunits where the LBD-S1 lobe in subunit B translates horizontally by 21.6 Å while the subunit C moves only by ∼9.5 Å. Further, the S2 lobes move towards center by 23.6 Å and 2.7 Å, respectively. As a result of this, the separation between S1 lobes in B and C reduces from 51.3 Å to 35.2 Å and that for S2 lobes reduces from 37.2 Å to 25.8 Å (Fig. 4f-h). The restricted movement of LBD for distal subunit C and larger movements of proximal subunit B leads to BC LBD dimer formation. We also compared the distal A/C and proximal B/D pairs (**Supplementary Fig. 11**) that show similar movements for ATD domains between 19.5 Å to 22.3 Å. The LBD domains of distal subunits A/C move by ∼ 9.7 Å while that for proximal B/D move by 21.6 Å for B and only 12.1 Å for subunit C, again highlighting the asymmetric arrangement of LBD in UBP310 bound state.

Owing to rearrangements at the LBD layer, there is also a substantial reorganization of LBD-TM linkers to go from desensitized to resting state. Consistent with 4-fold symmetric arrangement of LBDs in desensitized state, LBD-helix E separation in both distal A/C and proximal B/D subunit pairs is equal at ∼21 Å (**Supplementary Fig. 12 a, b**). Also helix-E is placed ∼4 Å lower in proximal B/D subunits when compared to distal A/C protomers (**Supplementary Fig. 12c**). In contrast, helix-E in UBP310 bound receptor has a separation of ∼22 Å for A/C pair similar to that in SYM complex but ∼45Å for the B/D subunits. Further, due to formation of BC dimer at the LBD layer, the helix-E in subunit B moves lower in vertical plane increasing the distance between helix-E from subunits B and C to ∼12 Å. Interestingly, the LBD helix-E from subunits A and D align in same plane in contrast to SYM complex where they are separated by ∼4 Å (**Supplementary Fig. 12**). These conformational changes in helix E-M3 are necessitated because of the asymmetric architecture of UBP310 bound receptor.

### N-glycans at the ATD-LBD interface

iGluRs are post-translationally modified with multiple N-linked glycans that are studded on both the extracellular ATD and LBD domains^17, 37^. Many of these glycans reside at the ATD-LBD interface and on the linkers connecting ATD and LBD (**Supplementary Fig. 1**) and have previously been shown to modulate gating properties^37, 38^ and trafficking of iGluRs^24, 39^ or interactions with other synaptic proteins^40^. In GluK3 desensitized EM map, we observed residual density at the ATD-LBD dimer interface, which was not satisfied by fitting of the ATD and LBD co-ordinates. Interestingly, the residual densities are in close proximity to potential N-linked glycosylation sites (Fig. 5a-b). In particular, the density near Asn 247 in ATD and Asn 402 in LBD appear more prominent in distal subunits as compared to proximal subunits (Fig. 5b). Similar extra density was also observed in the GluK2 (5KUF) EM map reported recently to a resolution of 3.8Å^31^. It is important to note that there is 2-fold symmetry of the ATD layer at the dimer of dimer interface and 4-fold symmetry at the LBD layer in a desensitized receptor. Due to this symmetry mismatch, the bottom of ATD and top of LBD domains of only the distal subunits are in close apposition and interact with each other unlike the ATD and LBD domains of proximal subunits. Further, the N-glycans are highly flexible molecules^41^ thus; they are likely to contribute poorly to EM density unless stabilized by glycan-glycan or glycan-protein interactions. Hence, the presence of this residual density at positions corresponding to N-glycan likely indicates protein-glycan interactions. It is interesting to note that N-glycans at Asn 247 and Asn 402 are suitably placed to mediate inter-domain interactions between ATD and LBD. N-glycan at Asn 247 from ATD seem to project down towards LBD and lies in close proximity with it; while glycan from LBD Asn 402 is oriented towards ATD and might interact with its lower lobe (Fig. 5b).

### Disrupting the N-glycan interactions affect GluK3 gating properties

To test whether these glycan interactions could affect gating properties of GluK3 receptors, we systematically mutated N-glycan sites; first individually and then in combination. Further, we carried out electrophysiological assays measuring key gating properties like rates of desensitization, recovery from desensitized state and EC_50_ values for agonist glutamate. The N-glycan knockouts at Asn 247, Asn 395 and Asn 402 were generated in wild-type GluK3 by mutating serine or threonine residues of the consensus N-glycosylation motif to alanine. On deletion of N-glycan, the rate of receptor desensitization was slowed down with a τ_des_ of 2.0 ± 0.3 ms (Asn 402); 2.1 ± 0.2 ms (Asn 247/Asn 402); 2.1 ± 0.2 ms (Asn 395/Asn 402) compared to wild type GluK3 that has a τ_des_ of 1.6 ± 0.1 ms. Similarly, the knockdown of Asn 247, Asn 395 glycan alone or in combination slowed the rate of receptor entry into desensitization with a measured τ_des_ of 1.8 ± 0.3 ms (Asn 247); τ_des_ of 1.6 ± 0.1 ms (Asn 395) and τ_des_ of 1.9 ± 0.2 ms, (Asn 247/Asn 395) respectively (Fig. 5c and f). Our results are consistent with those observed for GluK2 receptor triple N-glycosylation deletion at corresponding positions to GluK3 Asn 350, Asn 395 and Asn 402 and have also been shown to considerably slow down the rate of desensitization^37^.

Next, we tested whether these glycans affect the sensitivity of receptor for the agonists by measuring dose-response curves. Our results revealed that the dose response curves for GluK3 N-glycan mutants are left shifted with respect to wild type receptors. The macroscopic EC_50_ (glutamate) for wild-type receptors was calculated to be 9.5 mM while it was measured to be 7.2 mM (Asn 247); 3.7 mM (Asn 395); 6.5 mM (Asn 247/Asn 395); 5.3 mM (Asn 247/Asn 402) and 5.7 mM (Asn 395/Asn 402), respectively. While for single Asn 402 mutant it was calculated to be 10.3 mM, which is slightly higher than that of wild type GluK3 (Fig. 5d and g). Slower desensitization rate and lower EC_50_ values indicate that various N-glycan mutated receptors are more efficient and sensitive in responding to glutamate application. It’s interesting to note that the N-glycan knockouts for ATD and ATD-LBD linkers lowered glutamate EC_50_ while that on LBD (Asn 402) slightly increased it. However, all the three-glycan knockouts either individually or in combination slowed down desensitization rates.

### GluK3 N-glycan mutants recover faster from desensitized state

We next evaluated the recovery rates from desensitization by generating recovery currents via two applications of 30 mM glutamate at varying time intervals (two-pulse protocol) ranging from 50 ms to 5 s. We observe that all the GluK3 N-glycan mutants recovered from glutamate-evoked desensitized state faster than the wild type receptors with a τ_rec_ of 0.5 s (Asn 247); 0.8 s (Asn 395); 0.5 s (Asn 402) respectively for single knockouts and 0.6 s (Asn 247/Asn 395); 0.5 s (Asn 395/Asn 402) and 0.3s (Asn 247/Asn 402) for double glycan mutants. In contrast, the recovery rate of wild-type GluK3 had a τ_rec_ value of 0.9 s. Thus, the N-glycan mutants for positions 247, 402, 395/402 and 247/402 recover ∼1.7, 1.9, 1.9 and 3.2-fold faster than wild type receptors respectively, highlighting the importance of glycan at these positions in the receptor function (Table 2). In particular the glycans at position Asn 247 and Asn 402 alone or in combination seem to play key role in receptor desensitization and recovery. We hypothesized that glycans at these positions might affect receptor properties by mediating glycan-glycan and/or glycan-protein interactions. Hence, we next focused our attention to the putative N-glycan interacting residue on the receptor surface.

### Exploration of putative N-glycan interacting residues

It has been shown that the composition, content and length of N-glycans can be highly variable for iGluRs depending on the context of their expression. This in combination with flexibility of N-glycans makes it impossible to predict all the residues that might interact with them. Thus, we focused only on the residues that likely lay in close apposition to the N-acetyl glucosamine (NAG) residues of the oligo-mannose core at the ATD-LBD interface shown in Fig 5a-b. We mutated the potential interacting residues to their counterparts in GluA2 receptors. The core NAGs of Asn 402 N-glycan would likely lie in close proximity to ATD lower lobe (Fig. 5b). Interestingly, as in case of Asn 402 N-glycan knockout, single point mutants for positions Glu 239 and Tyr 243 to corresponding residues in GluA2 Leu (E239L) and Phe (Y243F) at the lower lobe of ATD slows down the receptor entry into desensitized state with a τ_des_ of 2.0 ±0.2 ms (E239L) and τ_des_ of 2.3 ±0.5 ms (Y243F) respectively (Fig. 6a). The mutant receptors also show faster recovery from desensitization than their wild-type counterparts with τ_rec_ of 0.3 s and 0.6 s for E239L and Y243F respectively similar to phenotype seen in case of N-glycan knockouts at Asn 402 (Fig. 6b). Interestingly, introducing a negatively charged residue by mutating arginine at position 242 to glutamate (R242E) in Y243F background (double mutant R242E/Y243F) showed a significant decrease in the rate of desensitization (τ_des_ of 3.3 ±0.5 ms) as in N-glycan knockouts but led to slower recovery from desensitized state (τ_rec_ of 4.0 s) in contrast to faster recovery rates observed for knockouts and other putative interacting residue mutants (Fig. 6c and d). The double mutant R242E/Y243F recovers ∼6.3 fold slower than Y243F mutant receptors and ∼4.2 fold slower than wild type GluK3. However, single mutation of R242E (corresponding residue in NMDA receptors) alters the measured receptor properties (τ_des_ and τ_rec_) τ_des_ of 2.1 ±0.1 ms and τ_rec_ of 0.8 s and is similar to wild type GluK3 (Fig. 6c-d). We don’t fully understand the reason for this observation but it’s likely due to modulation of protein-N-glycan interactions. Owing to 2-fold symmetry at ATD dimer and dimer of dimer interface, these mutations in a homotetrameric receptor in proximal subunits would lie close to the ATD dimer of dimer interface. However, these mutations are away from the central axis of receptor tetramer and are not likely to perturb tetrameric assembly. Hence, the affects seen are likely due to modulation of N-glycan-protein interactions on the distal subunits.

On evaluation of total surface expression of all the N-glycan and potential interacting residue mutants by surface biotinylation assay, it was found that all the mutants were able to reach the surface albeit in variable amounts. Surprisingly, N721A glycan knockout failed to reach cell surface (Fig. 6e; **Supplementary Fig. 13**). This N-glycan site lies at the LBD-TMD interface and was shown to be glycosylated in LBD expressed in insect cells^42^. Interestingly, it is conserved in all the kainate receptor subunits (GluK1-GluK5) but absent in NMDA and AMPA receptors. This suggests that glycosylation at this site might be important for assembly and trafficking of kainate receptors. However, it needs to be explored further to fully elucidate its role. Next, we investigated the potential interacting residues for Asn 247 N-glycan. The core NAGs of Asn 247 N-glycan likely lie in close proximity to S1 lobe of LBD. We checked double mutants Y744L/R745G and quadruple mutants Y744L/R745G/D746T/K747P for activity. However, in spite of reaching the cell surface (Fig. 6e; **Supplementary Fig. 13**) we could not record measurable currents from these receptors. Despite this, our glycan knockout assays highlight the importance of Asn 247 glycan in modulation of receptor functions.

To summarize, our structural and functional data suggest potential inter-domain interactions mediated by N-glycans, the perturbations of which alter receptor functions. It is noteworthy that the disruption or changes in interactions of N-glycans at position Asn 247 and Asn 402 that potentially mediate the ATD-LBD inter-domain interactions have maximum effect on the receptor properties. Our study highlights the contribution of N-glycans in enhancing the stability of the desensitized state and thus slowing down the recovery from desensitized state.

## Discussion

Kainate receptors are comparatively less studied than AMPARs and NMDA receptors but the current advancement in KAR knowledge indicates that they are involved in multifunctional neuronal activity and have a profound role in health and diseases^2, 43-45^. Using a multi-pronged approach, combining cryo-EM, X-Ray crystallography, electrophysiology, we studied GluK3 Kainate receptor structure in desensitized and closed state. Primarily, three major states exist in the gating cycle of glutamate receptor ion channels namely; resting, activated and desensitized state. Binding of agonist to the resting state receptor leads to a short-lived active state, which immediately relaxes to a desensitized state in order to relieve the strain caused by activated state onto linkers between LBD and TM^29, 34^. It quickly rearranges to a more energetically favorable desensitized conformation, where the LBD acquires the quasi 4-fold symmetry. Removal of agonist or binding of antagonists in *invitro* conditions allows the receptor to go in resting/closed state by rearranging LBD to a dimeric state, which leads to re-positioning of the linkers between LBD and TM^31^. It is well established that kainate receptors recover slower than AMPA receptors^46^. Also, the desensitized state in kainate receptors is ∼100 fold more stable than their AMPA counterparts^46^. N-glycans by virtue of their large size, position and chemical composition could mediate both intra and inter–domain interactions within the same subunit or other subunits in a tetramer. N-glycosylation corresponding to GluK3-Asn 402 in GluN1 (Asn 440) contributed to stability of closed clamshell LBD structure^47^. In this simulation study on isolated LBD, it was suggested that hydroxyl groups of N-glycan mannose are likely to interact with hydrophilic residues present on LBD only in closed conformation. While intra-domain LBD interactions were shown, no predictions for inter-domain interactions could be made as investigation was limited to isolated LBD domains and hence they cannot be precluded.

These interactions would potentially modulate receptor functions. Our GluK3-SYM and GluK3-UBP310 complex structures combined with electrophysiology based functional assays; identify structural elements on receptor surface, which potentially interact with N-glycans at Asn 247 and Asn 402 near the ATD-LBD interface (Fig. 7). These interactions would likely impede receptor recovery from desensitized state since the desensitized to resting state transition requires large-scale movements of the distal and proximal LBD domains in order to regain the dimeric configuration. The distal domains are stabilized in desensitized state likely by inter-domain interactions mediated by N-glycans at Asn 247 and Asn 402 apart from other protein-protein interactions. Our results provide a plausible explanation for the low potency of glutamate, which might be due to a partially recovered resting state in GluK3, reducing the efficacy of bound ligand in channel opening. This could also explain why the GluK3 receptors might desensitize faster even at sub-saturating glutamate concentrations. Disruption of the N-glycan mediated interactions likely allows full-recovery leading to slower desensitization, higher glutamate potency and faster recovery from desensitized state. Further, in the context of heteromeric receptors formed between GluK1-GluK3 receptors, our findings remain valid as both N-glycosylation sites and potential interacting residues are conserved (**Supplementary Fig. 1**). Similarly, for the receptors containing the “high-affinity” GluK4 and GluK5 subunits the residues corresponding to E239 and Y243 are aspartate and glutamate/aspartate respectively for the two sites in GluK4 and GluK5 subunits and may still interact with the N-glycan corresponding to Asn 402 on heteromeric assembly with GluK1-GluK3 subunits. Further, the interactions mediated by N-glycans would also affect the functional modulation by auxiliary proteins like Neto1 and Neto2 and has been shown earlier for GluK2^20^ but need more exploration in case of GluK3 receptors.

## Methods

### Construct design

We initially tried expression of the full-length rat GluK3 subunit containing point mutants R591Q (Q/R site) but this construct had very low expression and poor profile on size exclusion chromatography (SEC) and hence was not suitable for structural analysis. In order to achieve better surface expression, native signal peptide of GluK3 was replaced with that of GluK2 and was sub-cloned into the pEGBacMam vector^48^ for baculovirus based expression in mammalian cells using standard molecular biology techniques. To screen constructs via fluorescence detection^25^ and for affinity purification, a thrombin recognition site along with linker sequence (GLVPRGSAAAA) was inserted between GluK3 and the coding sequence for the A207K non-dimerizing EGFP mutant, with a C-terminal octa-histidine (His8) affinity tag. To improve solubility and stability, the GluK3 construct was truncated at C-ter Δ826 and cysteine residues at positions 86, 305 were mutated to threonine while cysteine 547 was replaced with valine. This construct had good expression and tetrameric receptor profile as screened via FSEC and hence was chosen for large-scale expression and purification. We refer to this construct as GluK3_EM_.

### Whole-cell voltage-clamp Recordings

In order to test the functionality of GluK3 wild type, GluK3_EM_ construct, and various mutants we performed whole-cell patch clamp recordings. Passage 20–35 human embryonic kidney 293 (HEK 293) cells were cultured on cover slips placed in a 35-mm polystyrene dish. Cells were transiently transfected either with GluK3_EM_ or co-tansfected with Wild type/mutant receptors along with GFP expressing plasmid (2 µg/dish) using Xfect reagent (Clontech). Currents were recorded from medium sized cells expressing a moderate level of fluorescence from either the fused EGFP in case of GluK3_EM_ or co-expressed EGFP and having a capacitance of ∼5-6 pF at 48-60 hours post transfection. Whole-cell patches were held under voltage clamp at −80 mV using an HEKA USB 10 amplifier. Ultrafast ligand application was achieved by using a two-barrel theta pipette ultra-fast perfusion system mounted on a piezoelectric device (Multichannel systems) controlled by Patchmaster software (HEKA). Responses were filtered at 3 kHz and digitized at 20 kHz. Recording pipettes were pulled (Sutter, P-1000) from borosilicate glass capillaries (1.5 OD × 1.17 × 100 L mm, Harvard Apparatus) and fire polished with an in-house developed microforge to reduce the tip diameter and get pipettes with a resistance between 2-3 Mega ohms. Pipettes were filled with intracellular solution (ICS) containing 30 mM CsCl, 100 mM CsF, 4 mM NaCl, 10 mM HEPES, 5 mM EGTA, 2 mM Na_2_ATP and 0.5 mM CaCl_2_, pH 7.2 and osmolarity ranging between 290-300 mosmol/L. Whole-cell patches were continually perfused with extracellular solutions (ECS) containing 150 mM NaCl, 2.8 mM KCl, 10 mM HEPES, and 0.5 mM CaCl_2_, pH 7.3 and osmolarity ranging between 295-305 mosmol/L at a flow rate of ∼0.2 ml/min. After attaining whole-cell voltage clamp, cells were raised off the dish, and positioned near the interface of the theta pipette solution streams. One stream contained control solution (ECS) while the other stream-contained ligands (_L_-glutamate, SYM or UBP310 at indicated concentrations) dissolved in the control solution. Longer applications of ligand i.e 100 ms were perfused to measure the whole-cell desensitization kinetics. The whole cell recordings were acquired using Patch master V2X90.2 (Heka Elektronick) 3 min after the formation of the whole-cell configuration. Raw data files were exported into Igor pro (ITX) and converted into abf files, compatible for pClamp by using ABF Utility. The macroscopic rate of desensitization was measured by the exponential fit to the decay of current from ∼90 % of its peak amplitude (*I*peak) to baseline and having a time constant of *τ*_des_. In presence of 30 mM glutamate the desensitization kinetics were fitted by using single exponential, 2-term fitting (Levenberg-Marquardt). For recovery from desensitization, a paired-pulse protocol was used and the peak amplitude of the second pulse was calculated as a percentage of total recovery. The time courses of recovery from desensitization were fit with a one-phase association exponential function. Values are reported as the mean ± SEM. Statistical tests were performed in Prism. Dose-response experiments were performed with different concentrations of glutamate in the range of 30 µM to 30 mM. Dose response curves were plotted as percentage of maximal response against Log [L-Glutamate concentrations and fitted with single exponential fits in Prism, version 8.0.1 (GraphPad Software, La Jolla, Ca, USA). Each data point indicates the mean of standard error and paired student T test was run for calculations of P values.

### Expression and Purification of GluK3 Receptor

GluK3_EM_ was transformed into DH10Bac cells to prepare bacmid for baculovirus generation following standard protocol^48^. HEK293 GnTI^-^ cells at density of 2.0-3.5 x10^6^ cells/ml were infected with P2 virus at a multiplicity of infection (MOI) value of ∼1. After 20 hours of incubation, 10 mM sodium butyrate was added to the culture flasks and transferred to 30°C for protein expression. Cells were harvested 70 hours post infection and stored at −80°C until further processing. Cell pellets were resuspended in ice-cold buffer (25–30 mL/L) containing 150 mM NaCl, 20 mM Tris (pH 8.0) and 1X Protease Inhibitors Cocktail (Roche). The resuspended cells were disrupted by ultrasonication (QSonica sonicator, 4 cycles of 90 sec (15 sec on/ 15 sec off) with power level 7 using medium size probe) with constant stirring. Care was taken to maintain the temperature of cell suspension below 12°C. The lysate was clarified by low-speed centrifugation and membranes were collected by ultracentrifugation (37,000 rpm, 60 min). Membrane pellets were homogenized, and solubilized for 45 min in buffer containing 150 mM NaCl, 20mM Tris (pH8.0), 29.5mM n-dodecyl-β-D-maltopyranoside, and 6 mM cholesterol hemisuccinate at 4°C. Insoluble material was removed by centrifugation (40,000 rpm, 60 mins) and cobalt-charged TALON metal affinity resin (∼7 mL) was added to the supernatant together with 10 mM imidazole to allow batch binding for 3 hours at 4°C. The beads were packed in the column, washed with 10 mM and 40 mM imidazole containing buffer (20 mM Tris, 150 mM NaCl, 0.75 mM n-dodecyl-β-D-maltopyranoside, 0.03 mM cholesterol hemisuccinate) until the OD at 280 nm reached close to zero and then bound receptors were eluted with buffer containing 250 mM imidazole. Peak fractions were pooled and concentrated to ∼1.9 mg/ml and digested overnight at 4 °C with thrombin at a 1:100 wt/wt ratio. GluK3 receptor tetramers were isolated by gel filtration chromatography (Superose 6 10/300) in a buffer containing 150 mM NaCl, 20 mM Tris (pH 8.0), 0.75 mM n-dodecyl-β-D-maltopyranoside, and 0.03 mM cholesterol hemisuccinate were concentrated to ∼1.7 mg/mL (MWCO 100 kDa). The final n-dodecyl-β-D-maltopyranoside and cholesterol hemisuccinate concentration calculated based on fold reduction of protein volume during concentration was ∼3.5 mM and 0.15 mM respectively.

In order to trap the purified receptors in different states, GluK3 receptor was incubated with ligands directly before imaging. 2S, 4R-4-methylglutamate (SYM) at a final concentration of 2mM was added to protein to trap desensitized state whereas, 100 µM UBP310 was used to stabilize the receptor in resting state. Screening of the concentration and effect of these agonist and antagonist onto stability of the receptor was checked by FSEC analysis (data not shown).

### Specimen vitrification and Cryo-electron microscopy

For both GluK3-SYM and UBP310 complex, Quantifoil R1.2/1.3 Au 300 mesh grids were glow discharged for 90 secs at 15mA. 2.5µl of protein at 1.7 mg/ml was applied to the grid thrice followed by blotting for 5 s at 100% humidity, 4°C and vitrified by plunging into ethane cooled by liquid nitrogen using vitrobot. The grids were clipped and loaded into a 300 kV Titan Krios microscope equipped with Falcon 3 direct-detector camera. Images were recorded in super-resolution counting mode with nominal magnification of ∼59,000X with the defocus range of −2.3 to −3.3 µm in steps of 0.3. Total 25 frames were collected in a movie with 60 s exposure, and a dose rate of 1.07 e/Å/s with a pixel size of 1.37Å for SYM and dose rate of 1.17 e/Å/s with a pixel size of 1.41Å for UBP310 complex respectively.

### Electron Microscopy Image Processing and model building

All the images (719 images for GluK3-SYM and 1693 for GluK3-UBP310) were subjected to beam induced drift correction using UCSF MotionCor2^49^ followed by CTF estimation via Gctf^50^. Manual curation was carried out to remove micrographs with large ice contamination or poor CTF fits. RELION 2.1^51^ was used to pick ∼1000 particles manually and subjected to reference-free 2D classification. Selected 2D classes from this step were used as reference for automated particle picking from entire datasets. Autopicked particle stacks were then exported to cryoSPARC^52^ for GluK3-SYM and cryoSPARC v2 for GluK3-UBP310 to carryout 2D classification. Data were cleaned up *via* iterative rounds of 2D classification and subsequently removal of classes with unclear features, ice contamination or carbon. Initially, 77276 particles selected for GluK3-SYM and 138369 particles for GluK3-UBP310. After successive rounds, 14648 and 54001 particles were selected for the GluK3-SYM and GluK3-UBP310 respectively. These were used to generate reference free *abinitio* 3D reconstruction. Additional round of 2D classification and removal of poorly defined particles selected 9730 particles that were utilized for 3D classification and refinement for SYM bound receptor in C1 symmetry. Homogenous refinement as implemented in cryoSPARC resulted into ∼7.4Å resolution map according to gold standard Fourier Shell Correlation (FSC) for GluK3-SYM complex. For GluK3-UBP310, 3D reconstruction and homogeneous refinement by imposing C1 symmetry lead to ∼8 Å resolution map. Sorted stack of 38853 particles was subjected to particle local motion correction followed with 2D classification and *abinitio* reconstruction into two 3D classes in cryoSPARC V2. The best 3D class (27194 particles) was further subjected to homogenous refinement resulting into a map with resolution of ∼8.3 Å. This was followed with non-uniform and local refinement that improved the map resolution to ∼ 8.1 Å and ∼7.7 Å (0.143 FSC) respectively. Final maps for both the complexes were sharpened and ResMap^53^ and Local Resolution module as implemented in cryoSPARC workflow was used for estimation of local resolution.

### Crystallization and Structure Determination of GluK3 LBD with SYM

GluK3 LBD S1S2 domain from residues N402-K515 of S1 domain linked by GT linker to P638-P775 of S2 domain (numbering according to mature polypeptide) was cloned into pET22b vector with a N-terminal hexa histidine tag (MHHHHHH) followed by LVPRGS thrombin cleavage site. Protein was expressed in *E. coli* Rosettagammi 2 (DE3) strain and purified by following the previously published protocol^12^. Purified and concentrated protein was dialyzed against agonist SYM containing buffer. The final protein at a concentration of ∼19 mg/ml in buffer containing 20 mM HEPES pH 7.0, 200 mM NaCl, 5mM Zinc acetate, 1 mM EDTA and 500 µM SYM was used for crystallization trials at 22°C. Best crystals grew in 50 mM HEPES, pH 7.0, 4% PEG 8000, 100 mM NaCl and were cryo-protected by transferring to solution containing 25 % glycerol as cryoprotectant and all the components of protein and crystallization mother liquor. X-ray diffraction data was collected at European Synchrotron Radiation Facility (ESRF), France. The data was indexed, scaled and merged using iMosflm^54^ and SCALA^55^. Finally, structure was solved by molecular replacement using crystal structure of LBD-Kainate complex (PDB: 3U92) as template with PHASER^56^ and refined using Phenix^57^ to a final resolution of 1.83 Å (**Supplementary Table 1**). Structural analysis and figures were prepared using UCSF Chimera^32^ and PyMOL^58^. Bound SYM was well resolved in our crystal structure alongwith interacting residues (**Supplementary Fig. 14, and 15**). Structure of GluK3-SYM LBD is very similar to GluK2-SYM structure reported earlier with r.m.s.d of 0.78 for superimposition of Cα atoms (**Supplementary Fig. 16**).

### Model building of the full-length receptor

Model for the GluK3-SYM tetramer was built by rigid body fitting in UCSF Chimera of 4 copies each of GluK3 amino terminal domain ATD (PDB code, 3OLZ and for ligand binding domain GluK3-LBD-SYM complex crystal structure determined in this study was used. Further, for GluK3 TM, sequence was threaded onto the transmembrane domain of GluK2 (PDB code: 5KUF) and was used for fitting into EM density. For GluK3-UBP310, PDB: 3OLZ used for ATD whereas LBD and TM were threaded onto LBD of 5CMK (LY466195 bound monomer) and TM of 5KUF respectivley. Coot^59^ was used to jiggle fit coordinates into EM density followed by multiple rounds of real space refinement (rigid body) of the complete model using PHENIX^33^. All the four subunits fit well into the EM map of both GluK3-SYM and GluK3-UBP310 shown in **Supplementary Fig. 7 and 8** with clear density of ATD, LBD and TM (pre-M1, M3 and M4) segments.

### Site Directed Mutagenesis

All the mutations were made in wild type GluK3 receptors following standard protocol for site directed mutagenesis and confirmed by sequencing of the entire GluK3 coding region. To disrupt N-linked glycosylation, S/T was substituted to alanine in N-linked consensus glycosylation sequences (NXS/T). Thus, T249, T397, S404 and T723 were mutated to alanine to disrupt N-linked glycosylation at N247, N395, N402 and N721 respectively. To check the putative N-glycan interacting residues for N247 glycans, residues Y744, R745, D746, K748 were mutated to (the corresponding residues in GluA2) Y744L/R745G (double mutant) and Y744L/R745G/D746T/K748P (quadruple mutant). Similarly, mutants E239L, R242E and Y243F were made to test potential N402 glycan interacting residues.

### Surface Biotinylation Assay

To evaluate surface expression of mutant receptors we carried out surface biotinylation assay. HEK293 cells were seeded in 6-well plates and transfected with various mutants along with wild type GluK3. After 40 hours of transfection, cells were washed with 1 ml ice-cold PBS / CaCl_2_ / MgCl_2_ twice. Following which, they were treated with 0.3 ml of 0.5 mg / ml Sulfo-NHS-SS-biotin solution for 30 min at 4 °C. Post incubation, biotin solution was removed and reaction was stopped by addition of biotin-quenching solution (50 mM glycine in PBS/CaCl_2_/MgCl_2_ buffer). This was followed by three rounds of washing with ice-cold PBS buffer. Cells were lysed in 100 µl of solubilization buffer (150 mM NaCl, 20 mM Tris, pH 8.0, 1mM PMSF), centrifuged to remove debris. Supernatant was mixed with 50µl of 50 % slurry of neutravidin agarose beads and rotated for 5 hours at 4 °C to allow binding of biotinylated proteins. Following this, the beads were washed extensively for removal of unbound proteins. Post final wash, supernatant was removed and 25µl of 4x SDS loading buffer was added to the beads and heated for 5 mins at 95 °C. Eluted proteins were resolved via SDS-PAGE, electroblotted onto PVDF membrane and probed with Anti-GluR6/7 monoclonal Antibody (Sigma) to identify surface expressed wild type and mutant GluK3 receptors. Blots were analyzed via ImageJ^60^ to quantitate total and surface expressed protein (Fig. 6d; **Supplementary Fig. 16**).

## Statistics

No statistical methods were used to predetermine sample size. The experiments were not randomised, and the investigators were not blinded to allocation during experiments and outcome assessment.

## Data availability

The cryo-EM density reconstruction and final model were deposited with the Electron Microscopy Data Base (accession code EMD-XXXX) and with the Protein Data Bank (accession code XXX). All other relevant data supporting the key findings of this study are available within the article and its <u>Supplementary Information</u> files or from the corresponding author upon request.

## Author Contributions

Jyoti Kumari optimized GluK3 construct, expressed and purified protein, carried out molecular biology, biochemical experiments and processed EM data with assistance from J.K., R.V. carried out the electrophysiology experiments. Janesh Kumar supervised the overall project design and its execution. All authors contributed to analysis and preparation of manuscript and approve the final draft.

## Acknowledgments

The research program in our laboratory is supported by the Wellcome Trust DBT India Alliance (IA/I/13/2/501023). Dr Kumar is an Intermediate Fellow of the Wellcome Trust/DBT India Alliance. Jyoti Kumari thanks University Grants Commission, India for senior research fellowship. R.V. is supported by SERB-N-PDF fellowship (N-PDF/2016/002621). Dr. Mark L. Mayer kindly gifted the various iGluR constructs that were subcloned and used for construct optimization and mutational studies. Dr. Eric Gouaux kindly provided the pEG BacMam vector. Access to EM was provided by the National Electron cryo-microscopy facility at the Bangalore Life Sciences Cluster. Funding for this was provided by a grant from the Department of Biotechnology, Government of India, DBT/PR12422/MED/31/287/2014. We thankfully acknowledge kind assistance of Dr. Vinothkumar Kutti Ragunath, NCBS, Bangalore in grid preparation and EM data collection.

## Corresponding author

Correspondence to Janesh Kumar

