## Supplementary Information for "Structural Insights into GluK3-kainate Receptor Desensitization and Recovery"

```
GluK3EM    1 MPHVIRIGGIFEYADGPNAQVMNAEEHAFRFSANIINRNRTLLPNTTLTYDIQRIHFHDSFEATKKACDQ  70
GluK3WT    1 MPHVIRIGGIFEYADGPNAQVMNAEEHAFRFSANIINRNRTLLPNTTLTYDIQRIHFHDSFEATKKACDQ  70
GluK2WT    1 TTHVLRFGGIFEYVE---SGPMGAEELAFRFAVNTINRNRTLLPNTTLTYDTQKINLYDSFEASKKACDQ  67
GluA2WT    1 SSNSIQIGGLFPRGA-------DQEYSAFRVGMVQFSTS-----EFRLTPHIDNLEVANSFAVTNAFCSQ  58

GluK3EM   71 LALGVVAIFGPSQGSTTNAVQSICNALEVPHIQLRWKHHPLDNKDTFYVNLYPDYASLSHAILDLVQSLK  140
GluK3WT   71 LALGVVAIFGPSQGSCTNAVQSICNALEVPHIQLRWKHHPLDNKDTFYVNLYPDYASLSHAILDLVQSLK  140
GluK2WT   68 LSLGVAAIFGPSHSSSANAVQSICNALGVPHIQTRWKHQVSDNKDSFYVSLYPDFSSLSRAILDLVQFFK  137
GluA2WT   59 FSRGVYAIFGFYDKKSVNTITSFCGTLHVSFITP---SFPTDGTHPFVIQMRPDLKG---ALLSLIEYYQ  122

GluK3EM  141 WRSATVVYDDSTGLIRLQELIMAPSRYNIRLKIRQLPIDSDDS-----RPLLKEMKRGREFRIIFDCSHT  205
GluK3WT  141 WRSATVVYDDSTGLIRLQELIMAPSRYNIRLKIRQLPIDSDDS-----RPLLKEMKRGREFRIIFDCSHT  205
GluK2WT  138 WKTVTVVYDDSTGLIRLQELIKAPSRYNLRLKIRQLPADTKDA-----KPLLKEMKRGKEFHVIFDCSHE  202
GluA2WT  123 WDKFAYLYDSDRGLSTLQAVLDSAAEKKWQVTAINVGNINNDKKDETYRSLFQDLELKKERRVILDCERD  192

GluK3EM  206 MAAQILKQAMAMGMMTEYYHFIFTTLDLYALDLEPYRYSGVNLTGFRILNVDNPHVSAIVEKWSMERLQA  275
GluK3WT  206 MAAQILKQAMAMGMMTEYYHFIFTTLDLYALDLEPYRYSGVNLTGFRILNVDNPHVSAIVEKWSMERLQA  275
GluK2WT  203 MAAGILKQALAMGMMTEYYHYIFTTLDLFALDVEPYRYSGVNMTGFRILNTENTQVSSIIEKWSMERLQA  272
GluA2WT  193 KVNDIVDQVITIGKHVKGYHYIIANLGFTDGDLLKIQFGGANVSGFQIVDYDDSLVSKFIERWSTLEEKE  262

GluK3EM  276 APRAESGLLDGVMMTDAALLYDAVHIVSVTYQRAPQMTVNS--------LQCHRHKAWRFGGRFMNFIKE  337
GluK3WT  276 APRAESGLLDGVMMTDAALLYDAVHIVSVCYQRAPQMTVNS--------LQCHRHKAWRFGGRFMNFIKE  337
GluK2WT  273 PPKPDSGLLDGFMTTDAALMYDAVHVVSVAVQQFPQMTVSS--------LQCNRHKPWRFGTRFMSLIKE  334
GluA2WT  263 YP----GAHTATIKYTSALTYDAVQVMTEAFRNLRKQRIEISRRGNAGDCLANPAVPWGQGVEIERALKQ  328

GluK3EM  338 AQWEGLTGRIVFNKTSGLRTDFDLDIISLKEDGLEKVGVWSPADGLNITEVAKGRGPNVTDSLTNRSLIV  407
GluK3WT  338 AQWEGLTGRIVFNKTSGLRTDFDLDIISLKEDGLEKVGVWSPADGLNITEVAKGRGPNVTDSLTNRSLIV  407
GluK2WT  335 AHWEGLTGRITFNKTNGLRTDFDLDVISLKEEGLEKIGTWDPASGLNMTESQKGKPANITDSLSNRSLIV  404
GluA2WT  329 VQVEGLSGNIKFDQN-GKRINYTINIMELKTNGPRKIGYWSEVDKMVVTLTELPSG-NDTSGLENKTVVV  396

GluK3EM  408 TTLLEEPFVMFRKSDRTLYGNDRFEGYCIDLLKELAHILGFSYEIRLVEDGKYGAQDDK-GQWNGMVKEL  476
GluK3WT  408 TTLLEEPFVMFRKSDRTLYGNDRFEGYCIDLLKELAHILGFSYEIRLVEDGKYGAQDDK-GQWNGMVKEL  476
GluK2WT  405 TTILEEPYVLFKKSDKPLYGNDRFEGYCIDLLRELSTILGFTYEIRLVEDGKYGAQDDVNGQWNGMVREL  474
GluA2WT  397 TTILESPYVMMKKNHEMLEGNERYEGYCVDLAAEIAKHCGFKYKLTIVGDGKYGARDADTKIWNGMVGEL  466

GluK3EM  477 IDHKADLAVAPLTITHVREKAIDFSKPFMTLGVSILYRKPNGTNPSVFSFLNPLSPDIWMYVLLAYLGVS  546
GluK3WT  477 IDHKADLAVAPLTITHVREKAIDFSKPFMTLGVSILYRKPNGTNPSVFSFLNPLSPDIWMYVLLAYLGVS  546
GluK2WT  475 IDHKADLAVAPLAITYVREKVIDFSKPFMTLGISILYRKPNGTNPGVFSFLNPLSPDIWMYVLLACLGVS  544
GluA2WT  467 VYGKADIAIAPLTITLVREEVIDFSKPFMSLGISIMIKKPQKSKPGVFSFLDPLAYEIWMCIVFAYIGVS  536

GluK3EM  547 VVLFVIARFSPYEWYDAHPCN---PGSEVVENNFTLLNSFWFGMGSLMQQGSELMPKALSTRIIGGIWWF  613
GluK3WT  547 CVLFVIARFSPYEWYDAHPCN---PGSEVVENNFTLLNSFWFGMGSLMQQGSELMPKALSTRIIGGIWWF  613
GluK2WT  545 CVLFVIARFSPYEWYNPHPCN---PDSDVVENNFTLLNSFWFGVGALMQQGSELMPKALSTRIVGGIWWF  611
GluA2WT  537 VVLFLVSRFSPYEWHTEEFEDGRETQSSESTNEFGIFNSLWFSLGAFMQQGCDISPRSLSGRIVGGVWWF  606

GluK3EM  614 FTLIIISSYTANLAAFLTVERMESPIDSADDLAKQTKIEYGAVKDGATMTFFKKSKISTFEKMWAFMSSK  683
GluK3WT  614 FTLIIISSYTANLAAFLTVERMESPIDSADDLAKQTKIEYGAVKDGATMTFFKKSKISTFEKMWAFMSSK  683
GluK2WT  612 FTLIIISSYTANLAAFLTVERMESPIDSADDLAKQTKIEYGAVEDGATMTFFKKSKISTYDKMWAFMSSR  681
GluA2WT  607 FTLIIISSYTANLAAFLTVERMVSPIESAEDLSKQTEIAYGTLDSGSTKEFFRRSKIAVFDKMWTYMRSA  676

GluK3EM  684 P-SALVKNNEEGIQRTLTAD--YALLMESTTIEYITQRN-CNLTQIGGLIDSKGYGIGTPMGSPYRDKIT  749
GluK3WT  684 P-SALVKNNEEGIQRTLTAD--YALLMESTTIEYITQRN-CNLTQIGGLIDSKGYGIGTPMGSPYRDKIT  749
GluK2WT  682 RQSVLVKSNEEGIQRVLTSD--YAFLMESTTIEFVTQRN-CNLTQIGGLIDSKGYGVGTPMGSPYRDKIT  748
GluA2WT  677 EPSVFVRTTAEGVARVRKSKGKYAYLLESTMNEYIEQRKPCDTMKVGGNLDSKGYGIATPKGSSLGTPVN  746

GluK3EM  750 IAILQLQEEDKLHIMKEKWWRGSG-CPEEEN---KEASALGIQKIGGIFI--------------------  795
GluK3WT  750 IAILQLQEEDKLHIMKEKWWRGSG-CPEEEN---KEASALGIQKIGGIFIVLAAGLVLSVLVAVGEFIYK  815
GluK2WT  749 IAILQLQEEGKLHMMKEKWWRGNG-CPEEES---KEASALGVQNIGGIFIVLAAGLVLSVFVAVGEFLYK  814
GluA2WT  747 LAVLKLSEQGVLDKLKNKWWYDKGECGAKDSGSKEKTSALSLSNVAGVFYILVGGLGLAMLVALIEFCYK  816

GluK3EM  795 ----------------------------------------------------------------------  795
GluK3WT  816 LRKTAEREQRSFCSTVADEIRFSLTCQRRLKHKPQPPMMVKTDAVINMHTFNDRRLPGKDSMSCSTSLAP  885
GluK2WT  815 SKKNAQLEKRSFCSAMVEELRMSLKCQRRLKHKPQAPVIVKTEEVINMHTFNDRRLPGKETMA-------  877
GluA2WT  817 SRAEAKRMKVAKNPQNINPSSSQNSQNFATYKEGYNVYGIESVKI-------------------------  861

GluK3EM  795 ---  795
GluK3WT  886 VFP  888
GluK2WT  877 ---  877
GluA2WT  861 ---  861
```

**Supplementary Fig. 1** Amino acids sequence alignment of rat GluK3$_{EM}$, GluK3WT, GluK2WT and GluA2WT. All the sequences are numbered as in mature polypeptide. Identical and conserved residues are shaded in grey color. Disordered region which correspond to no or weak density are highlighted in red. Cysteine mutations (C86T, C305T and C547V) are indicated in purple color. Potential N-linked glycosylation sites that were mutated are (N247, N395, N402 and N721) shown in green color. Potential N-glycan interacting amino acids that were mutated and tested are highlighted in blue.

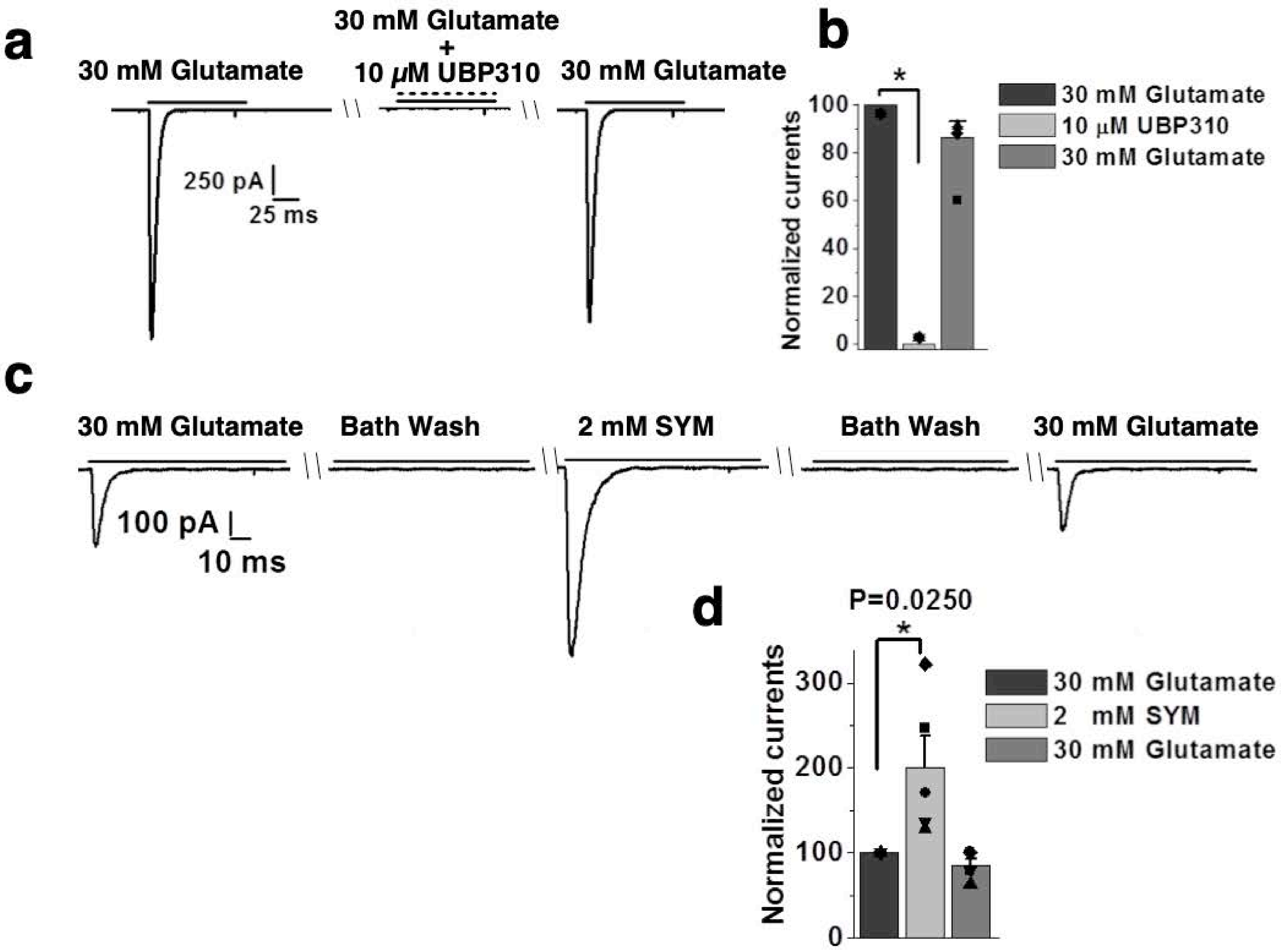

**Supplementary Fig. 2 a.** GluK3$_{EM}$ is blocked by UBP310 and potentiated by SYM application. **a**. Representative whole-cell traces for currents evoked by GluK3$_{EM}$ recombinantly expressed in HEK293T cells in response to application of 30 mM glutamate. Glutamate currents are blocked by co-application of 10 µM UBP310 **b**. shows normalized current amplitude for same (n=5). Panel **c** shows currents evoked in response to 30 mM glutamate followed with washing and successive application of 2 mM SYM. **d.** shows normalized current amplitudes for the same (n=6).

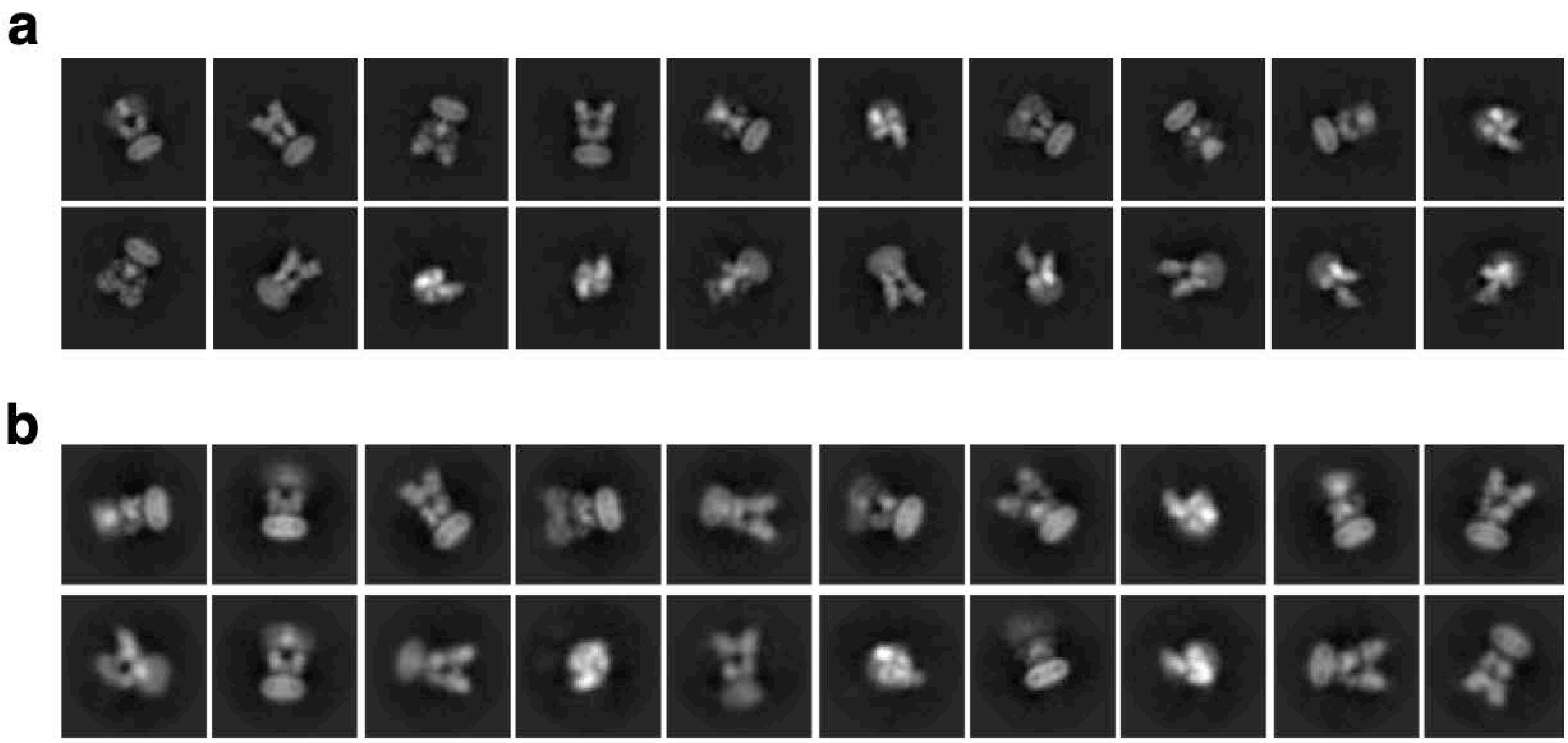

**Supplementary Fig. 3** 2D classification. Representative 2D classes are shown for **a.** GluK3-SYM and **b**. GluK3-UBP310 complexes.

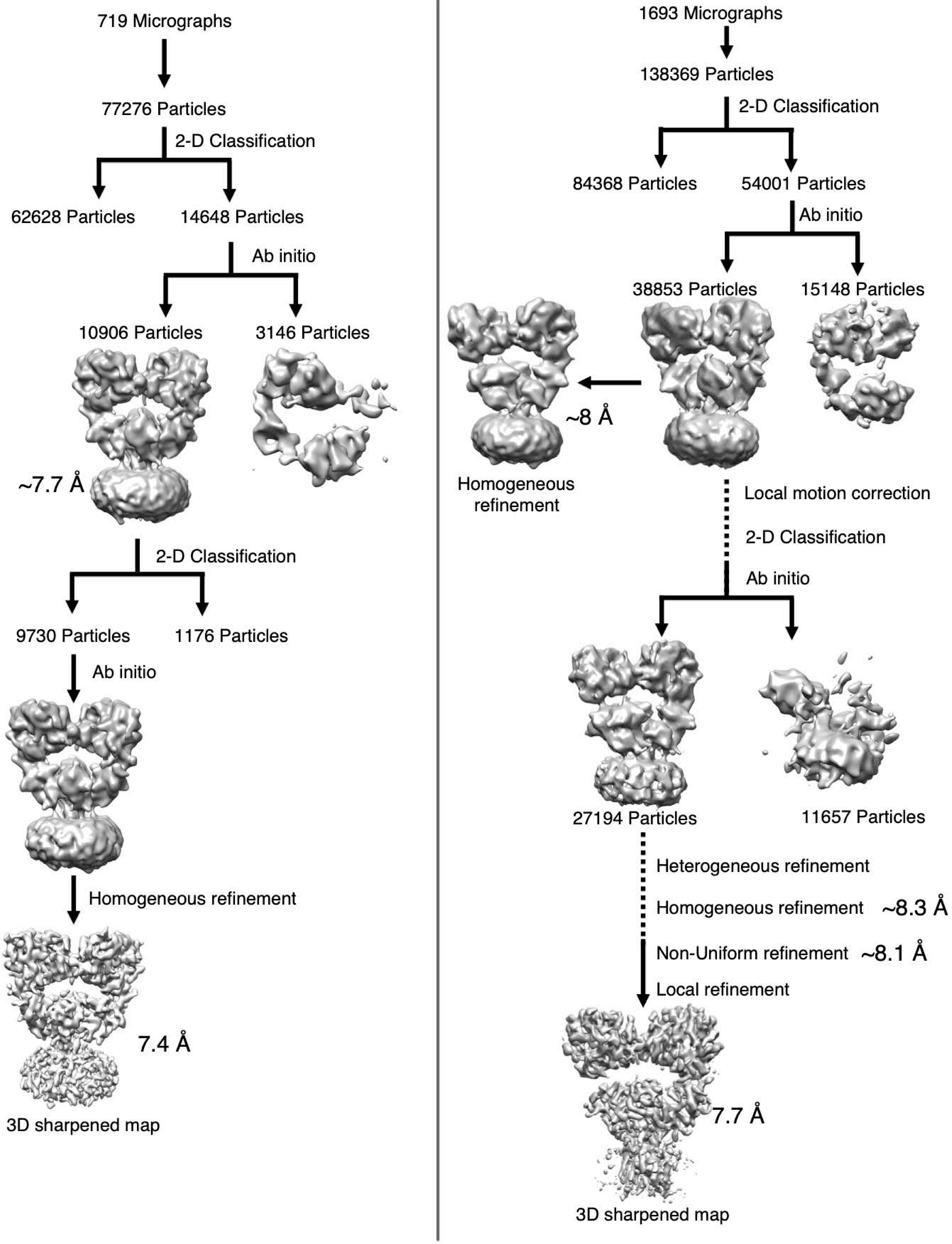

**Supplementary Fig. 4** Cryo-EM data processing work flow for **a.** GluK3-SYM and **b.** GluK3-UBP310.

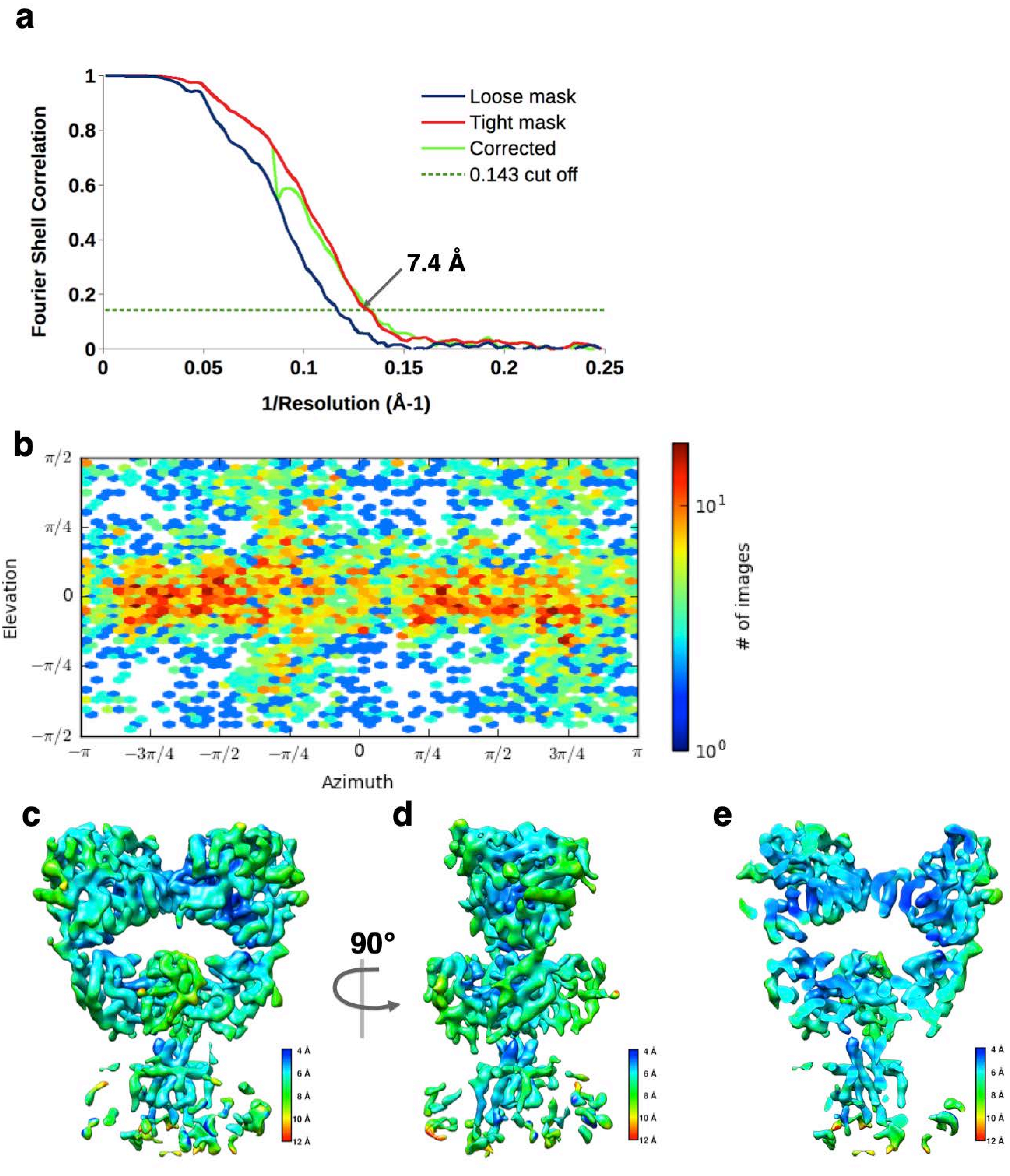

**Supplementary Fig. 5** 3D reconstruction for GluK3-SYM. **a.** FSC curves for the density map of SYM-bound GluK3 receptors showing 7.4 Å resolution as per Gold standard 0.143 FSC. **b** viewing direction distribution of particles used for 3D reconstruction over azimuth and elevation angles. **c.** shows the final density map colored according to local resolution estimated by RESMAP.

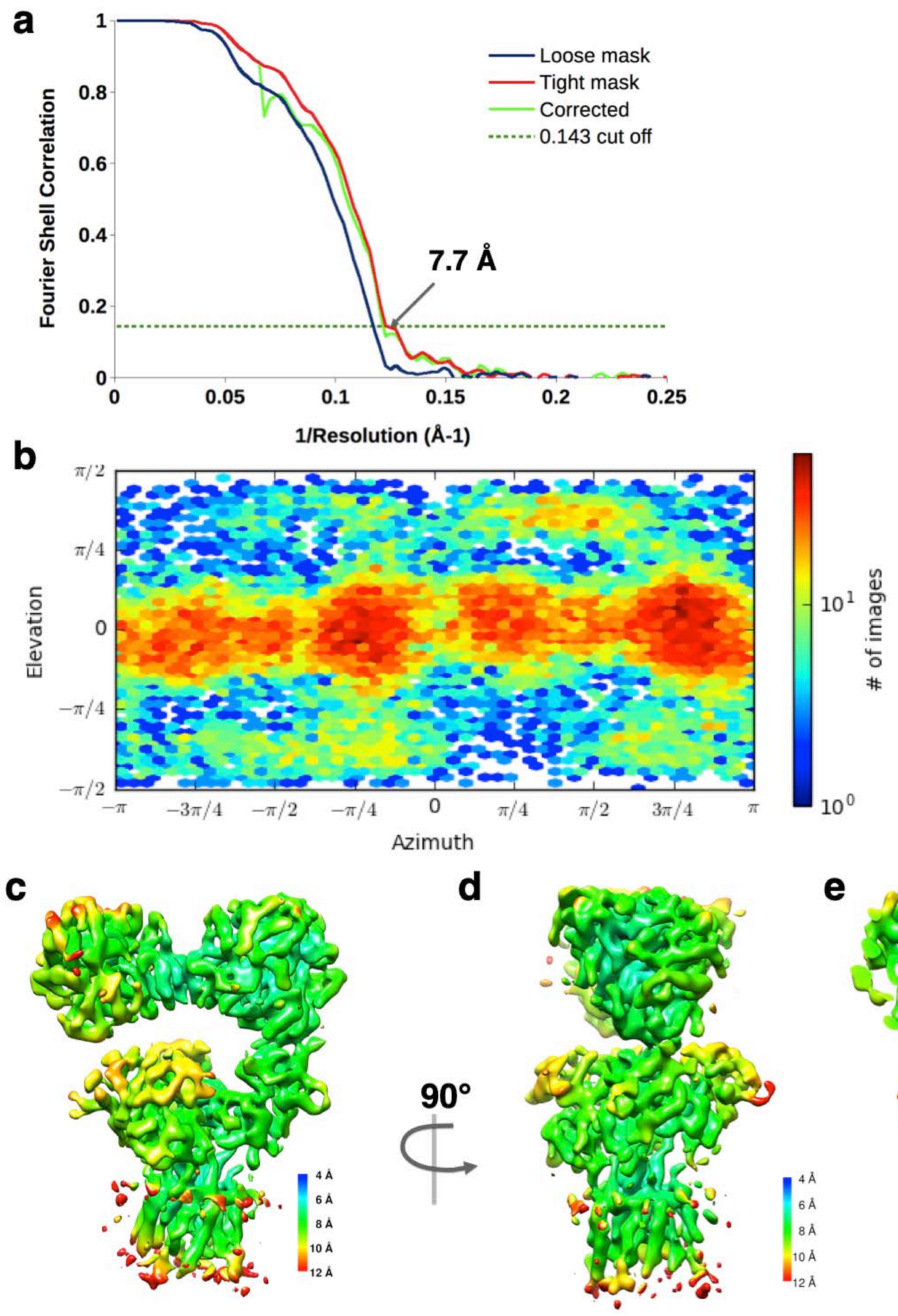

**Supplementary Fig. 6** 3D reconstruction for GluK3-UBP310. **a.** FSC curves for the density map of UBP310-bound GluK3 receptors showing 7.4 Å resolution as per Gold standard 0.143 FSC. **b.** viewing direction distribution of particles used for 3D reconstruction over azimuth and elevation angles. **c.** shows the final density map colored according to local resolution estimated by RESMAP.

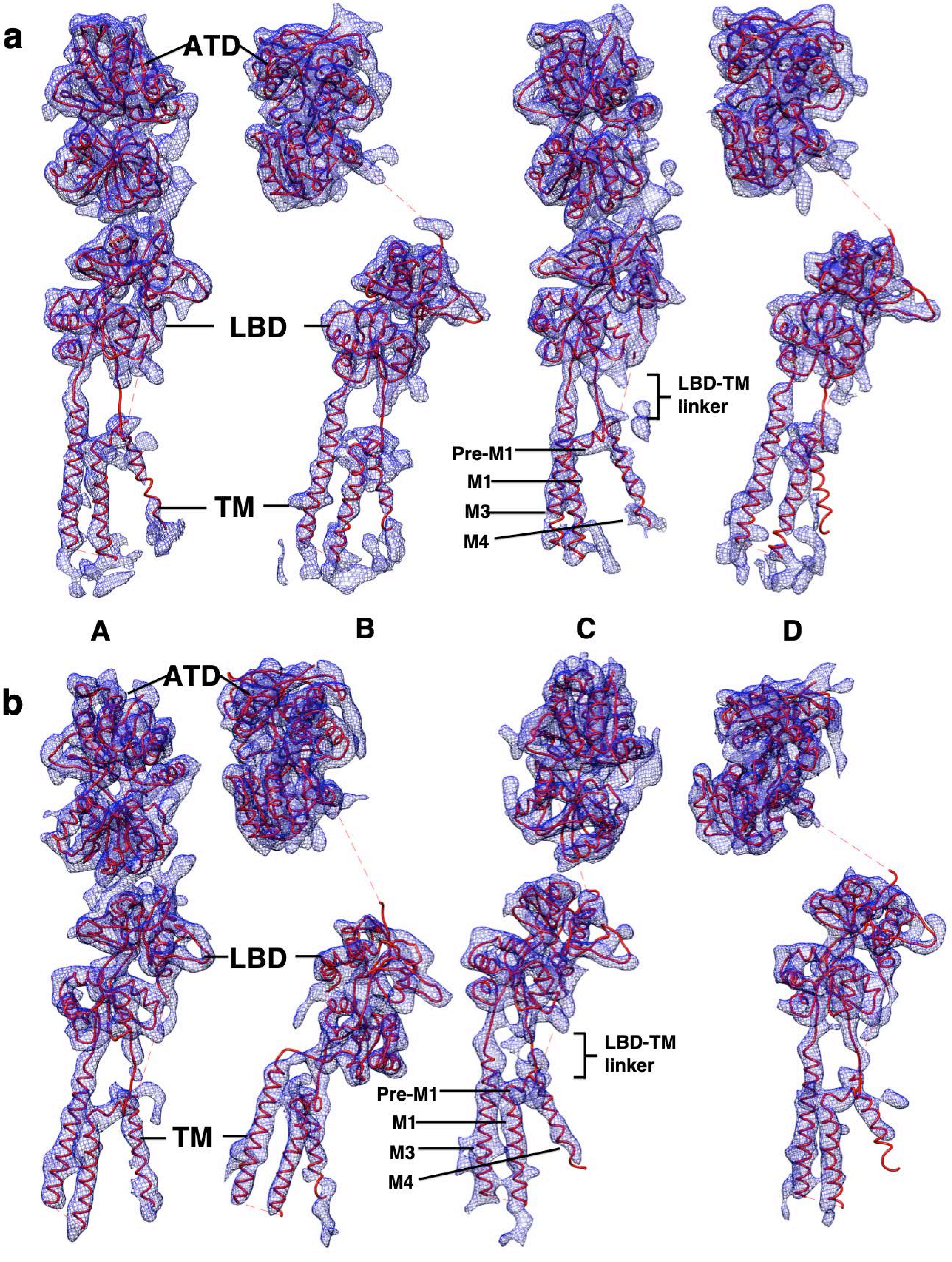

**Supplementary Fig. 7**. Segmented density map. Panels a and b represent segmented density map fitted with receptor protomers A-D represented in red ribbons shows the quality of density map for GluK3-SYM (**a**) and GluK3-UBP310 (**b**) respectively. Individual subunits are labelled as in receptor tetramer.

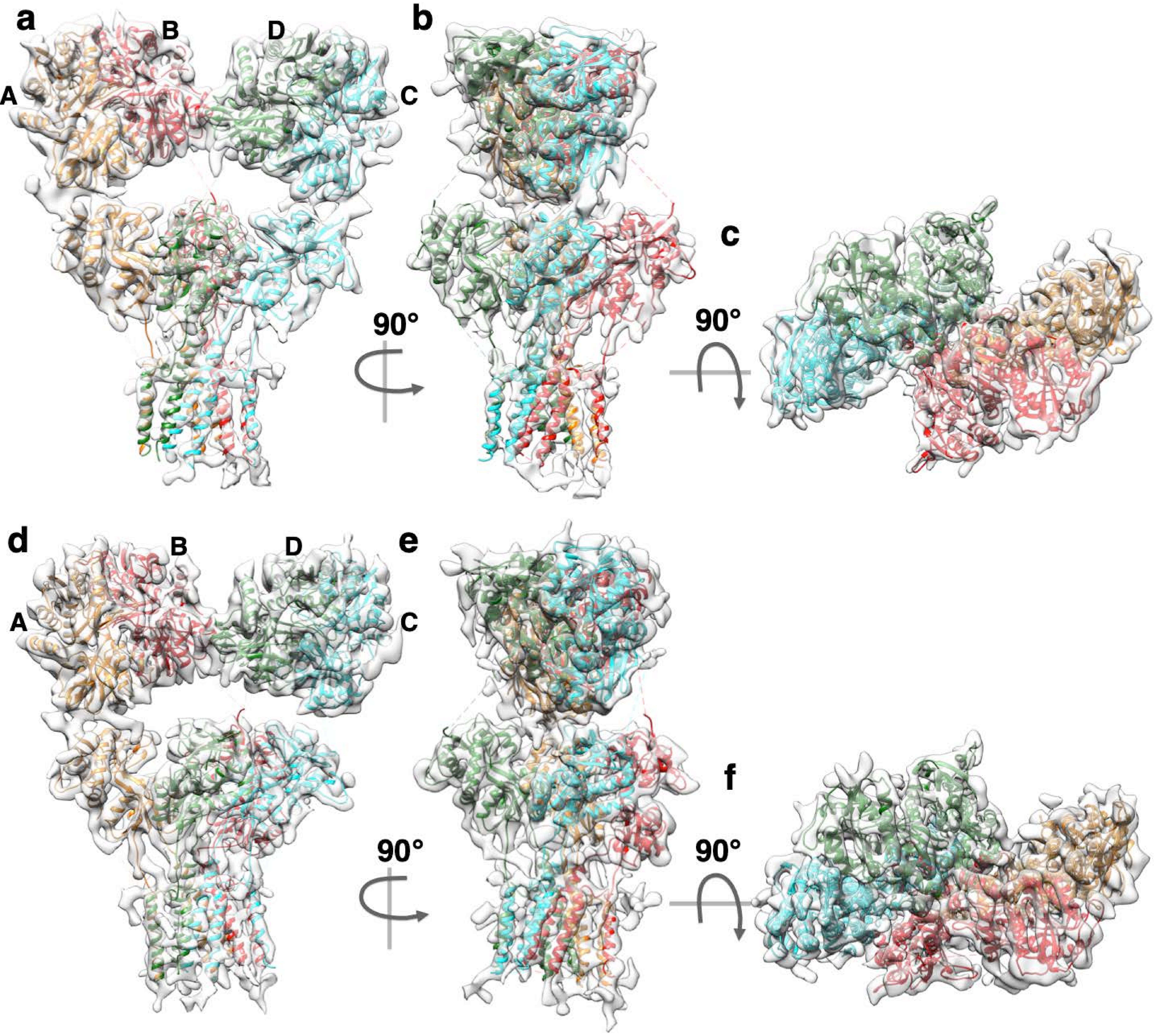

**Supplementary Fig. 8** Atomic models with each chain uniquely colored and fitted into EM density map for GluK3 desensitized (**a-c**) and closed/resting states (**d-f**). Panel **a.** and d show view parallel to membrane; **b** and **e** are 90° rotated views along Y-axis for **a** and **d** . Panels **c** and **f** show top views for **b** and **e** generated via rotation along X-axis.

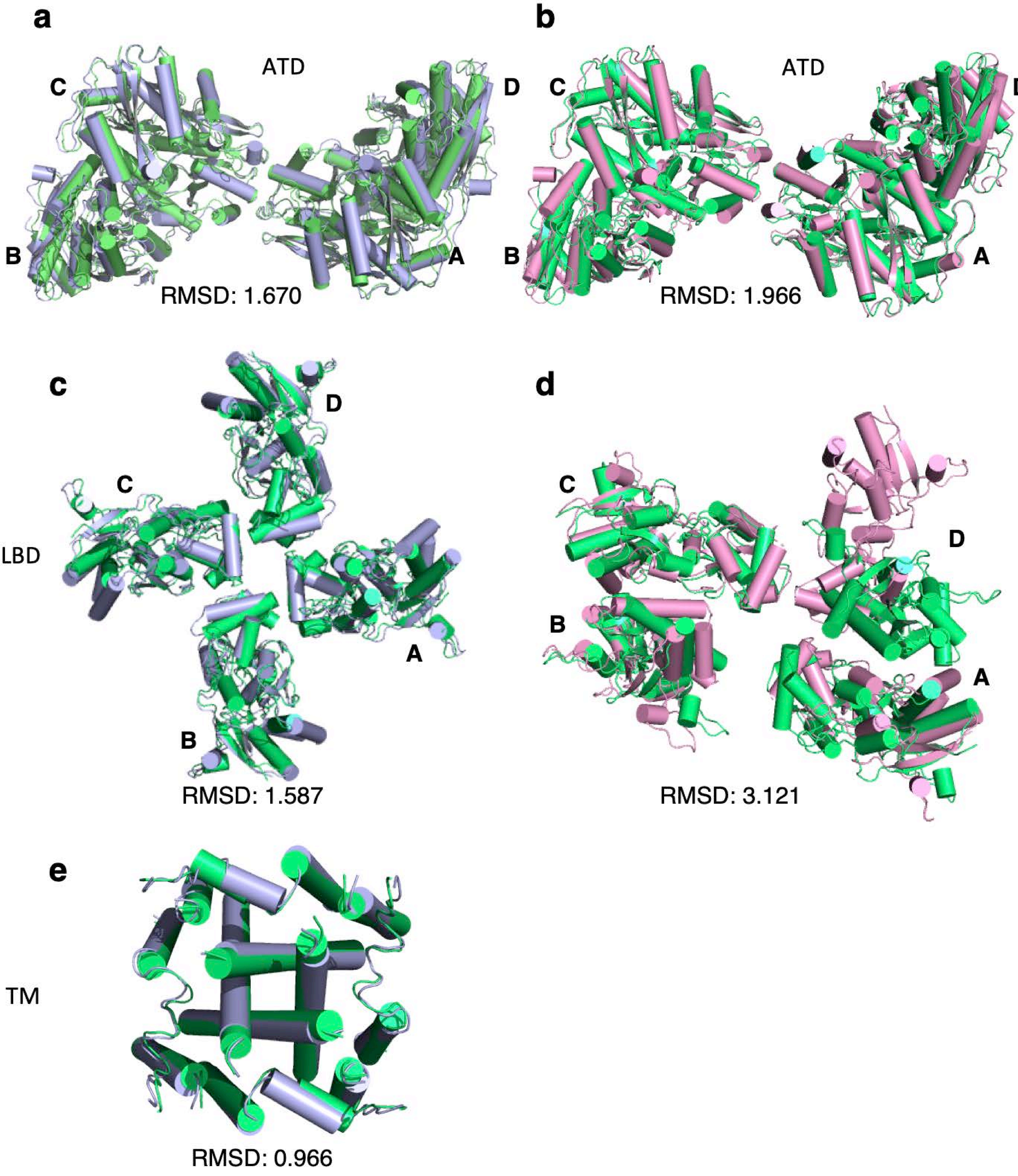

**Supplementary Fig. 9 Superimposition and comparison of GluK3-SYM and GluK3-UBP310 with GluK2-SYM (5KUF) and GluK2-LY(5KUH).** Individual domains of GluK2 structures were superimposed with that from GluK3 and shown as top views. Panels (**a, c, and e**) are superimposition (Cα atoms) of GluK3-SYM (blue) with GluK2-SYM (green) for ATD, LBD and TM domains. Panels (**b and d**) show superimpositions for GluK3-UBP310 (magenta) ATD and LBD domains with GluK2-LY (green) ATD and LBD.

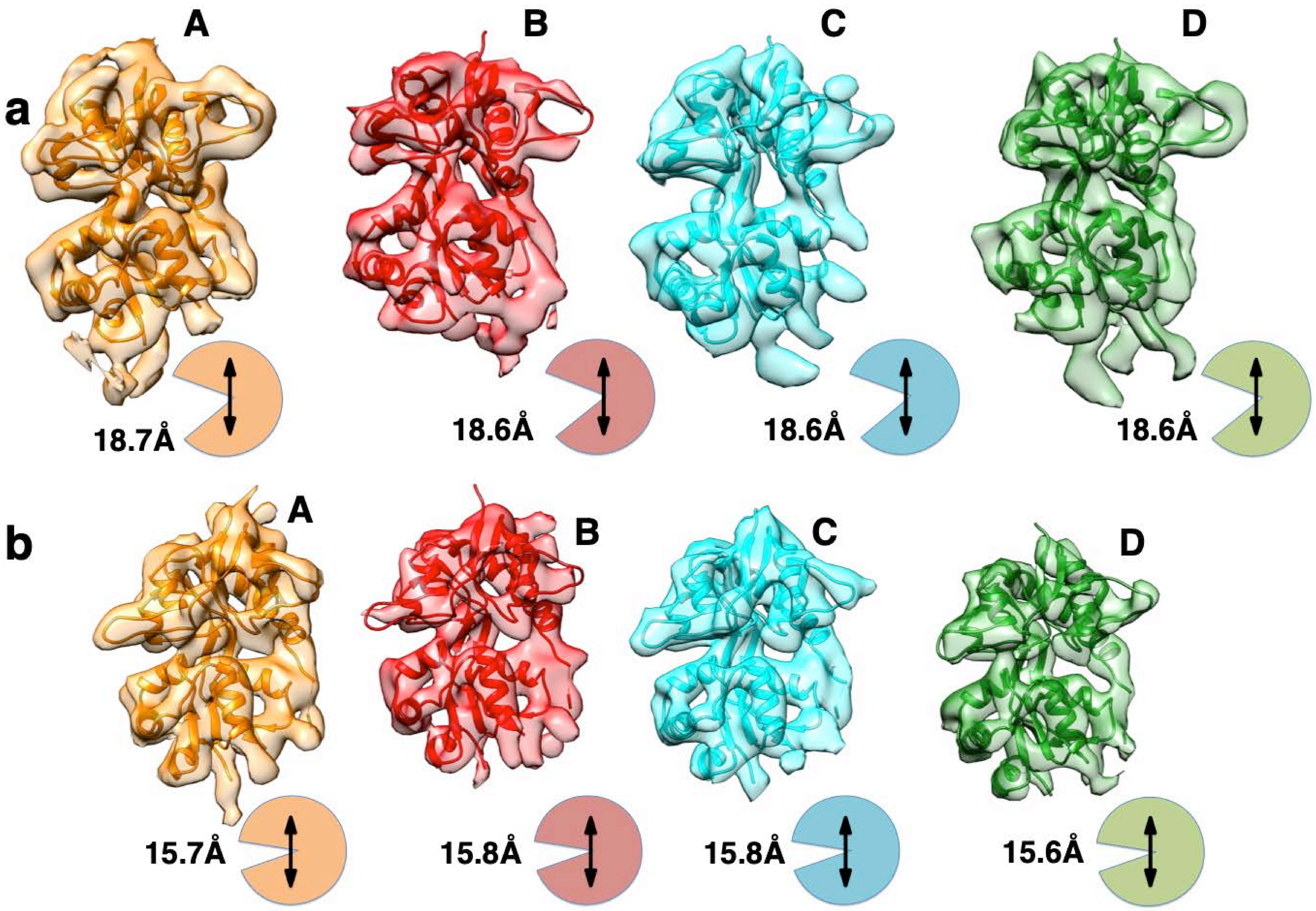

**Supplementary Fig. 10** Segmented EM map for LBD domain fitted with UBP310 bound (**a**) and SYM bound (**b**). Different subunits are represented in as orange-A, red-B, cyan-C, green-D respectively. Antagonist (UBP310) bound form shows an extended LBD cleft compared to agonist (SYM) bound form. Distance between Centre of mass (COM) for S1 and S2 lobes for the UBP310 bound form is ~18.6 Å compared to ~15.7 Å for SYM bound LBD consistent with the extended cleft in antagonist bound state. Cartoon representation of each LBD subunit is shown next to it with measured distances between COMs of S1 and S2 lobes.

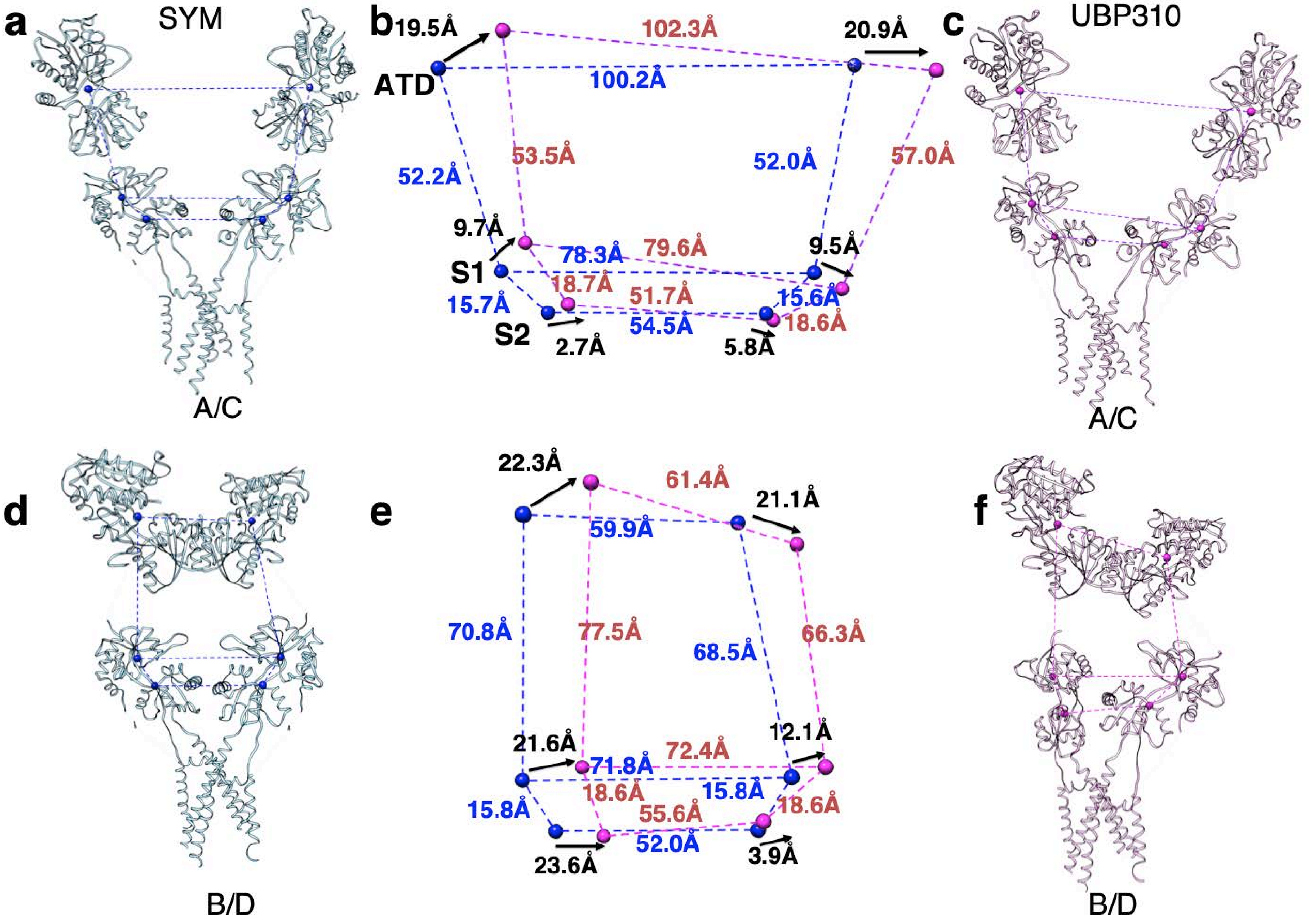

**Supplementary Fig. 11 Comparison between GluK3-SYM and GluK3-UBP310 during the transition from desensitized to resting state**. Arrangement of individual subunits for distal A/C and proximal B/D subunits are shown in panel (**a and d**) for GluK3-SYM and in (**c, f**) for GluK3-UBP310. Distances from the center of mass (spheres) between subunits for the ATD (top) and LBD S1 & S2 lobes are measured and indicated by dashed lines. Structures were superimposed using main-chain atoms of the TM domains. **b & e** show superimpositions of the parallelograms formed for the A/C (**b**) and B/D (**e**) subunits. The movements of the COM for domains between SYM and UBP310 structures are measured and indicated in black fonts, while the distances within structures are indicated in blue (SYM) and red (UBP310).

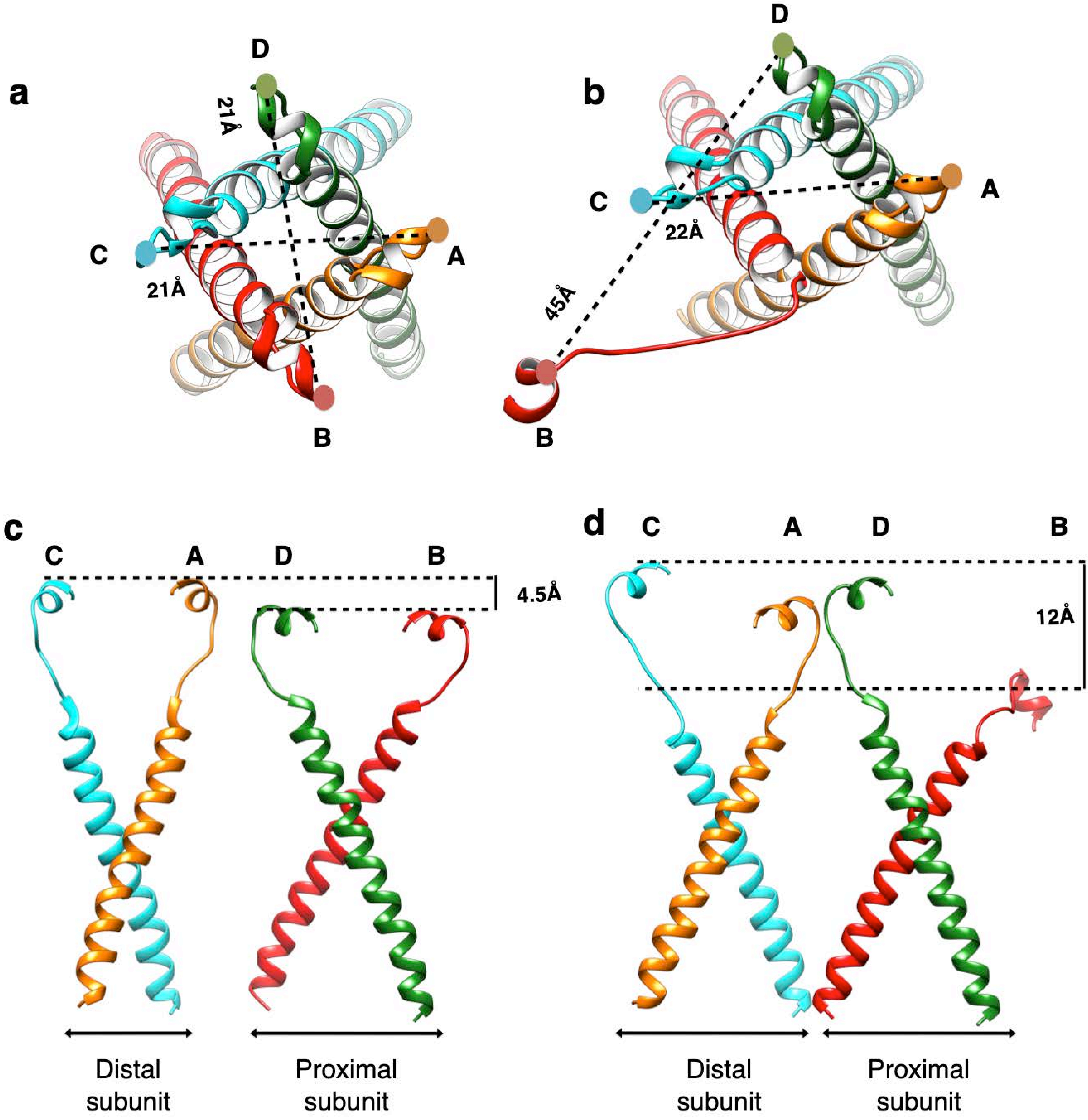

**Supplementary Fig. 12 Conformational changes at the M3, M3-S2 linkers and E helices for A/C and B/D subunits in GluK3-SYM (a, c) and GluK3-UBP310 (b,d) structures.** Top views of the LBD S2- helix E-M3 linker regioins for full-length receptors in complex with SYM (**a**) and UBP310 **(b**), measured distances between COMs of helix E for A/C and B/D pairs are shown. Panel **c** and **d** show the front views of the helix E-M3 segments for distal A/C and proximal B/D subunits to highlight the height and pitch differences between SYM and UBP310 bound states respectivley.

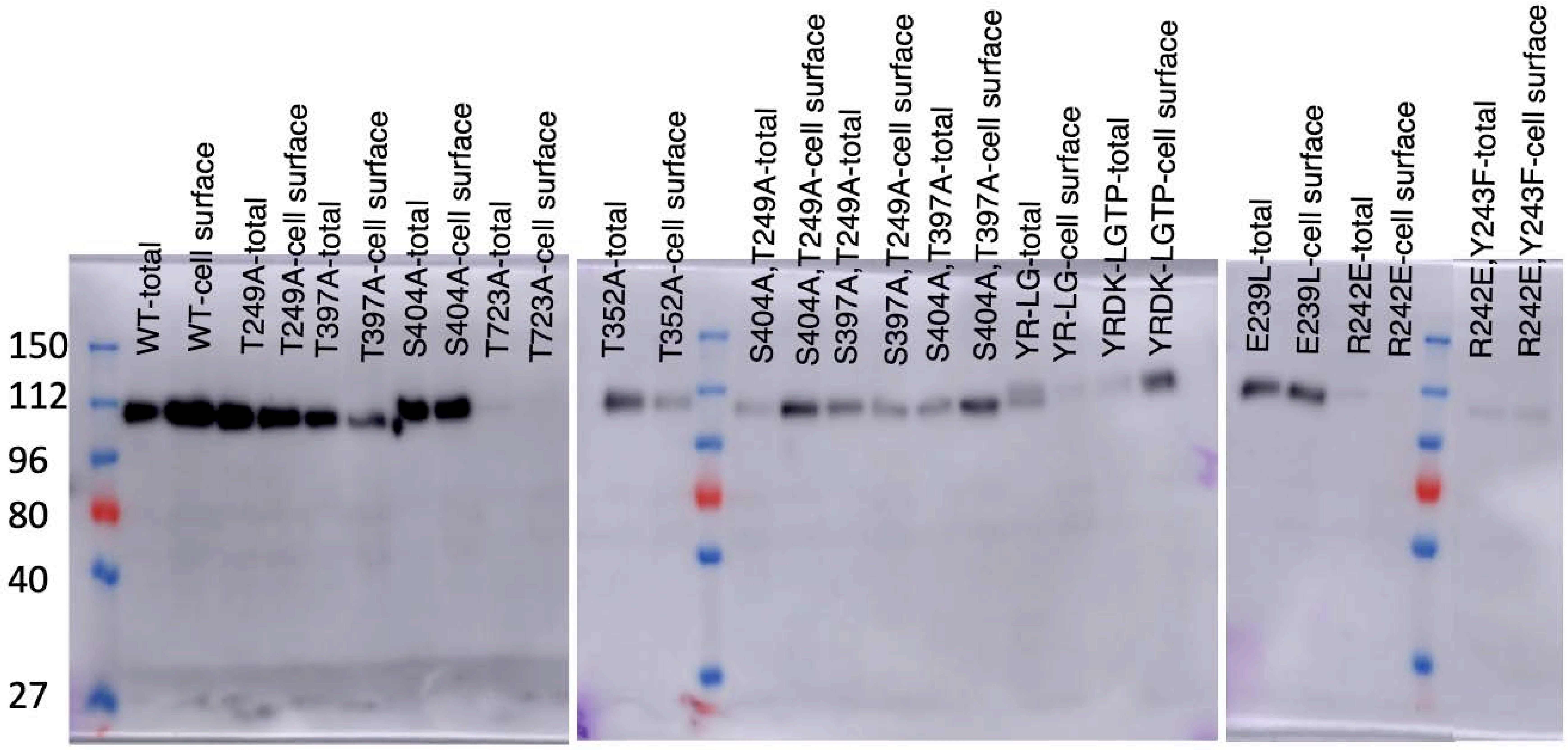

**Supplementary Fig. 13 Surface expression of GluK3 constructs**. Raw western blots are shown for evaluating surface expression of GluK3 constructs by surface biotinylation assay. Total and surface expression are shown. Blots were probed by immunoblotting with GluR6/7 monoclonal antibody (Sigma). Quantitation and analysis was carried out via ImageJ.

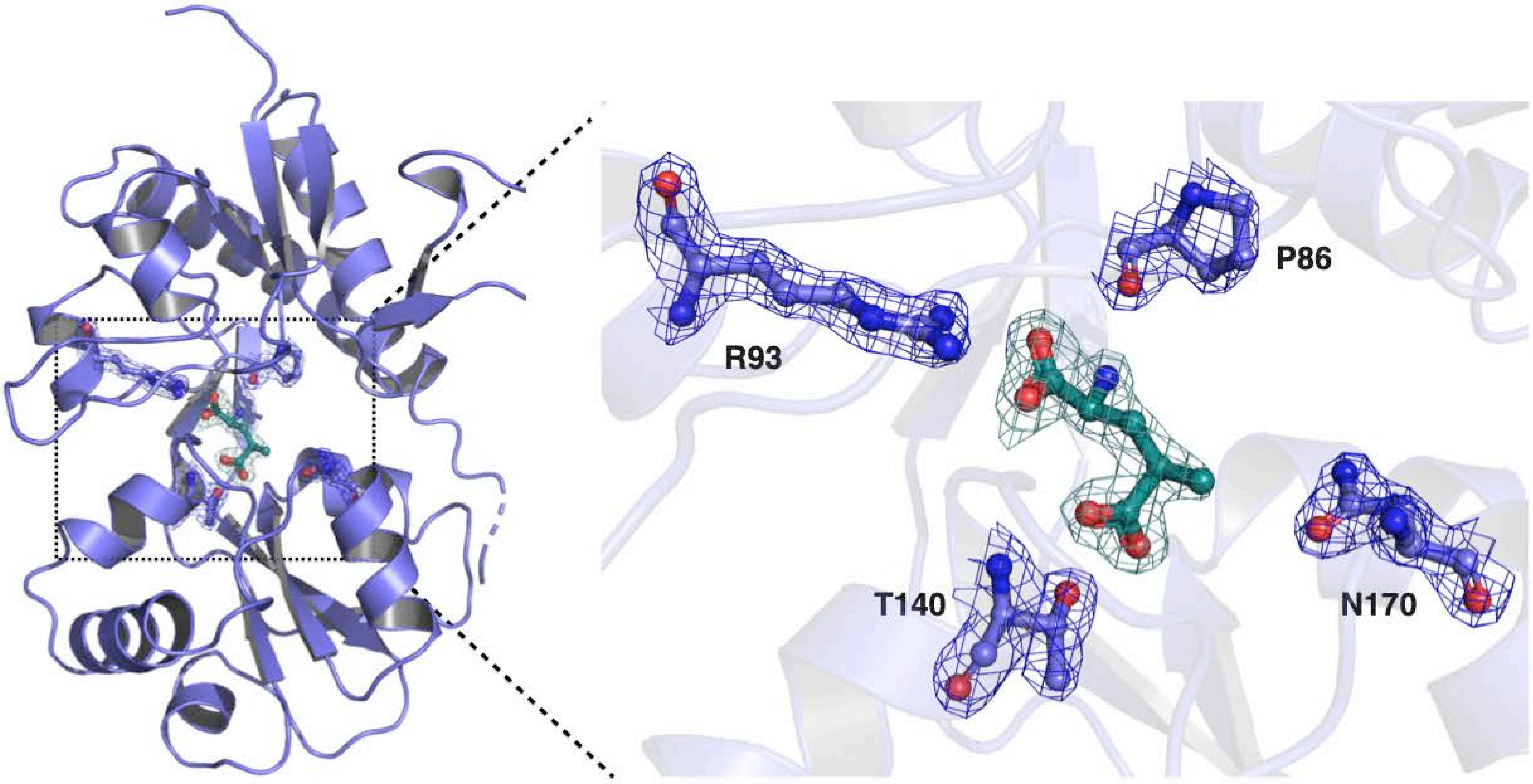

**Supplementary Fig. 14 Crystal structure of GluK3 LBD in complex with SYM**. **a**) 2Fo-Fc electron density map carved at 2σ. SYM and interacting residues are shown. **b** shows enlarged view centered at ligand showing the density for SYM and interacting residues Pro 86, Arg 93, Thr 140 and Asn 170.

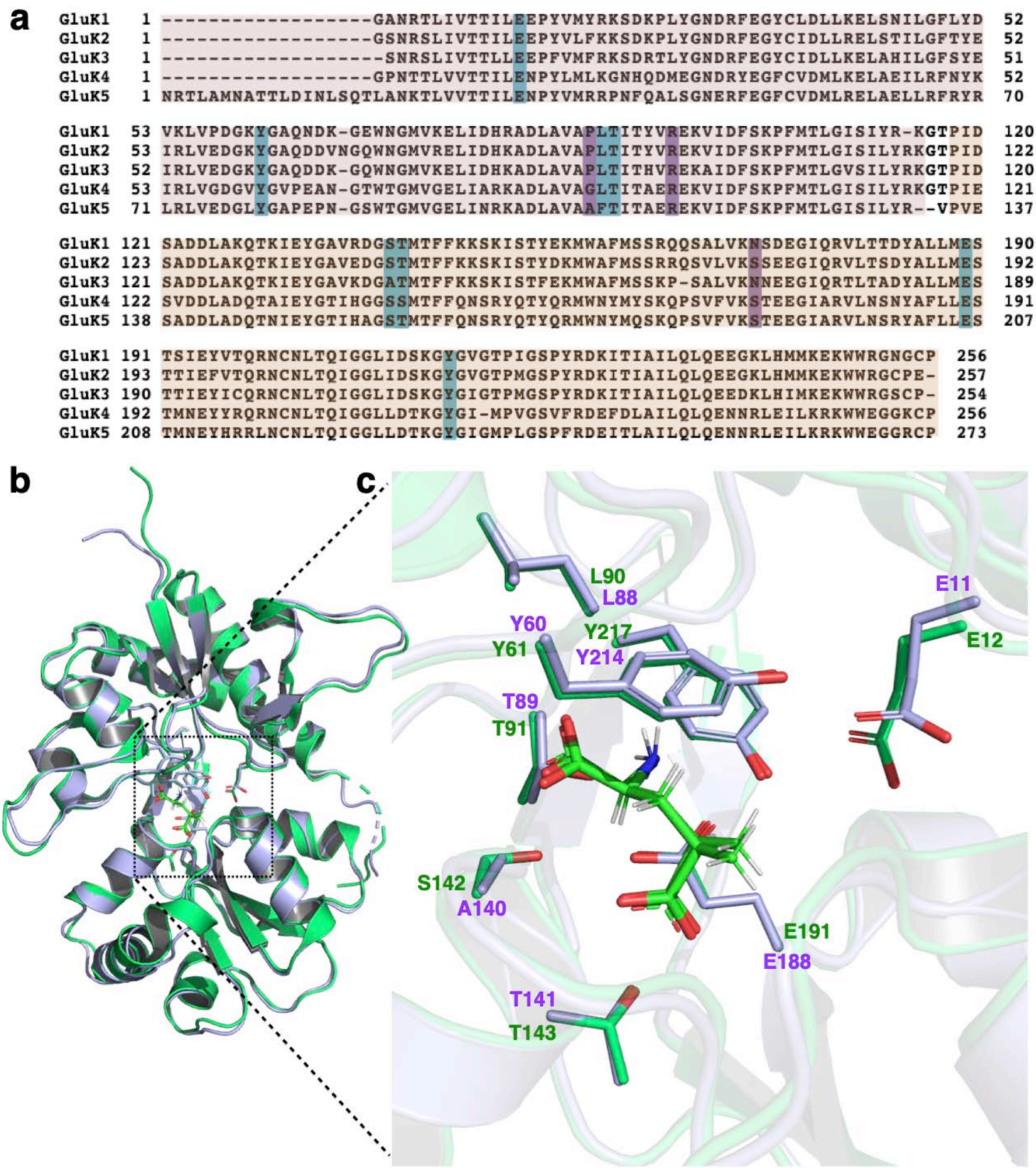

**Supplementary Fig. 15 Crystal structure of GluK3 LBD in complex with SYM**. **a**) Alignment of amino acids sequence for the S1 and S2 lobes of GluK1 (5M2V), GluK2 (5CMM), GluK3 (this study), GluK4 (5IKV) and GluK5 (uniprot) is shown. S1 lobe is shaded in light red while S2 lobe sequences are shaded in light orange color. Ligand binding residues are shaded in cyan. Panel (**b**) shows single S1S2 clamshell of GluK3-SYM (blue) superimposed with GluK2-SYM (5CMM, green) using C-alpha residues had an r.m.s.d of 0.476 indicating very similar structures. Panel (**c**) shows larger view of the binding pocket, all the interacting residues with SYM are conserved between GluK3 and GluK2 except one S142 which is a mutation of alanine to serine in GluK2 (5CMM).

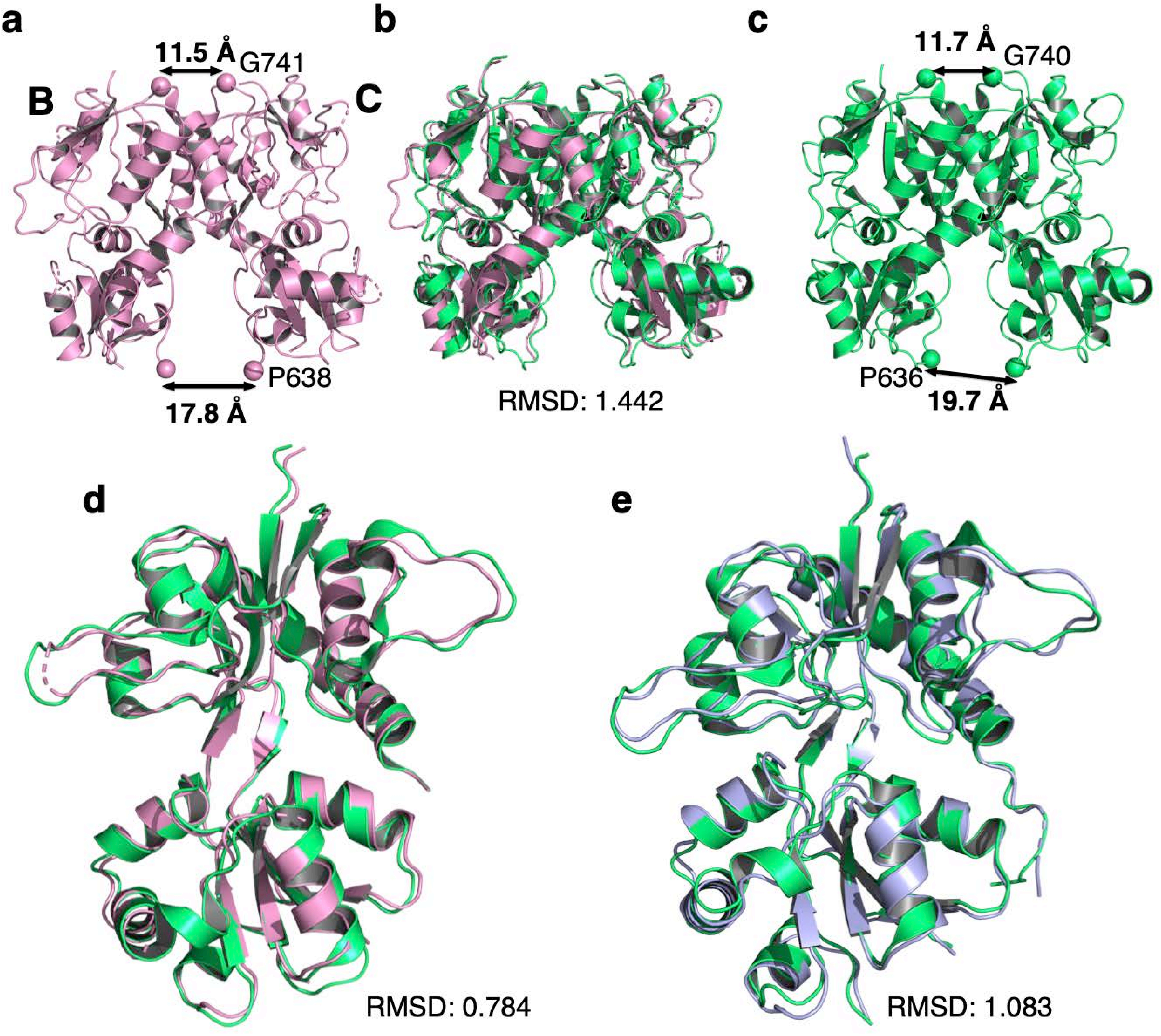

**Supplementary Fig. 16**. **UBP310 and SYM bound ligand binding domains**. GluK3-UBP310 LBD dimer (subunits BC) is depicted in pink ribbons in **a** is superimposed in **b** with LBD dimer from GluK2-LY(5KUH) (**c**) shown in green. Distances between Cα (shown as sphere) atoms at top (P638 and P636) and base (G741 and G740) of LBD dimer is shown indicating similar dimeric conformation. Superimposition of monomers for GluK3-UBP310 and GluK2-LY is shown in **d**. while LBD monomers from GluK3-SYM (blue) and GluK2-SYM (green) is shown in e. R.M.S.D values for each comparison is indicated.

**Supplementary Table 1** Crystallographic data collection and refinement statistics

| | **GluK3 (LBD) in complex with 2S,4R-4-methylglutamate** |
|---|---|
| **Data collection** | |
| Space group | P 2 21 2 |
| Cell dimensions a, b, c, (Å) | 56.46, 88.03, 130.47 |
| Cell angles α, β, γ, (°) | 90, 90, 90 |
| Wavelength (Å) | 0.97242 |
| Resolution range (Å) | 56.46 - 1.83 (1.89 - 1.83) |
| Completeness | 0.99 |
| Multiplicity | 5.1 |
| I/σI | 7.9 (1.3) |
| $R_{mease}$ (%) | 0.095 (0.979) |
| $CC_{1/2}$ (%) | 0.99 (0.11) |
| Wilson B-factor | 28.72 |
| **Refinement** | |
| No. of reflections | 57703 (5702) |
| $R_{work}/R_{free}$ (%) | 0.1985/ 0.2344 (0.4008/ 0.4478) |
| **Number of non-hydrogen atoms** | 4456 |
| Protein | 4061 |
| Ligands | 71 |
| **Average B-factor (A²)** | 46.84 |
| Protein | 46.53 |
| Ligands | 51.97 |
| solvent | 49.61 |
| **R.m.s. deviations** | |
| Bond Lengths (Å) | 0.013 |
| Bond angles (°) | 1.14 |
| **Ramachandran plot** | |
| Favored (%) | 97 |
| Allowed (%) | 3.4 |
| Disallowed (%) | 0 |
| Rotamer outliers (%) | 3.2 |
| *Highest resolution shell in parentheses | |
| 5% of reflections were used for calculation of $R_{free}$ | |
